# *Aedes albopictus* bionomics in Procida Island, a promising Mediterranean site for the assessment of innovative and community-based integrated pest management methods

**DOI:** 10.1101/2021.02.19.431954

**Authors:** B. Caputo, G. Langella, V. Petrella, C. Virgillito, M. Manica, F. Filipponi, M. Varone, P. Primo, A. Puggioli, R. Bellini, C. D’Antonio, L. Iesu, L. Tullo, C. Rizzo, A. Longobardi, G. Sollazzo, M. M. Perrotta, M. Fabozzi, F. Palmieri, G Saccone, R. Rosà, A. della Torre, M. Salvemini

**Affiliations:** Department of Public Health and Infectious Diseases, University of Rome La Sapienza, Italy; Department of Agriculture, University of Naples Federico II, Italy; Department of Biology, University of Naples Federico II, Italy; Department of Biodiversity and Molecular Ecology, Edmund Mach Foundation, San Michele all’Adige, Italy; Centro Agricoltura Ambiente “Giorgio Nicoli”, Crevalcore, Italy; Ministry of Education, University and Research, Italy; Centre Agriculture Food Environment, University of Trento, San Michele all’Adige (TN), Italy

## Abstract

The colonization of Mediterranean Europe and of other temperate regions by *Aedes albopictus* created in the last decades an unprecedented nuisance problem in highly infested areas, as well as a new public health threat due to the species competence to transmit exotic arboviruses, such as dengue, chikungunya and zika. The Sterile Insect Technique (SIT) and the Incompatible Insect Technique (IIT) are insecticide-free mosquito-control methods relying on mass release of irradiated/manipulated males which are believed to have a potential in complementing existing and only partially effective control tools. Testing and implementing these approaches are challenging and selection of study sites is an instrumental and crucial step. We carried out a 4-year study in Procida Island (Gulf of Naples, Italy) in strict collaboration with local administrators and citizens to estimate: i) the temporal dynamics, spatial distribution, and population size of *Ae. albopictus*; and ii) the dispersal and survival of irradiated males. Overall, results provide insights on the bionomics of the mosquito in Southern Europe and draw attention to Procida Island as an ideal site to test innovative control programs against *Ae. albopictus* which may be used in other Mediterranean and temperate areas.”

## Introduction

The Asian tiger mosquito *Aedes albopictus* (Skuse) (Diptera: Culicidae) is an invasive species that in the last few decades has greatly expanded its range from Southeast Asia to all other continents except Antarctica, mostly thanks to passive transportations of its drought-resistant eggs via used tire trade^1^. Since the first identification in Albania in 1979^2^, this exotic species has extended its distribution in several European countries thanks to its ability to adapt to seasonal variations and to use various man-made water containers for oviposition. In Italy, one of the most heavily infested country in Europe, *Ae. albopictus* was first reported in 1990 in Genova (north-west Liguria region)^3^ and quickly spread over the whole territory, in particular in the North-eastern area and central and southern coastal areas, including major and minor islands^4,5^. *Ae. albopictus* represents a relevant public health risk due to its vector competence for pathogens such as arboviruses and filarial worms^6^. In European Mediterranean regions it has been involved in the last decade in various autochthonous transmission events of chikungunya virus (CHKV) (France^7^ and Spain^8^), dengue virus (DENV) (Croatia^9^ and France^10^) and, more recently, of Zika virus (ZIKV) (France^11^). In Italy, *Ae. albopictus* caused in 2007 the first European chikungunya outbreak, with >200 human cases in North East Italy (Emilia Romagna)^12,13^. Ten years later, a second chikungunya outbreak occurred in Central and South Italy (Lazio and Calabria), with >500 cases, including cases in the metropolitan city of Rome^14,15^ and in a coastal village of South Italy (Calabria)^14^.

As for many other insect pests, mosquito spreading is contrasted to date mainly using chemical pesticides. The negative impact of the use/overuse of these compounds on the environment and on human health, the problem of growing insecticide resistance in mosquito populations, including *Ae. albopictus*^16–18^ and the absence of available vaccines against most arboviruses, render the development of eco-sustainable alternative mosquito-control methods an urgent need, as recently underlined by WHO^19^.

Promising complementary methods for mosquito control are represented by the Sterile Insect Technique (SIT)^20^ and the Incompatible Insect Technique (IIT)^21^. SIT relies on mass rearing and mass release of radiation-sterilized males into target areas to suppress local infesting insect populations. SIT technique has been successfully applied in the frame of area-wide integrated pest management (AW-IPM) programs^22^, against several insect species including agricultural pests (*Ceratitis capitata*, *Cydia pomonella*), livestock pests (*Cochliomyia hominivorax*) and vector species such as the tsetse fly *Glossina austeni*^23–26^. SIT potential against mosquitoes has been demonstrated for the first time in Italy with a three-year long study based on the release of 2 million sterile males of *Ae. albopictus*. By releasing 896-1,590 sterile males/ha/week in small villages it was possible to induce egg sterility in the range 18.72 – 68.46 % causing a significant decline in the egg density^27–29^. Starting from this encouraging pilot studies, the development of SIT against mosquitoes has rapidly advanced in recent years thanks to the research and coordinating efforts of the Joint Food and Agriculture Organization of the United Nations (FAO)/International Atomic Energy Agency (IAEA) Insect Pest Control Subprogramme and their collaborators, involved in the development of mass rearing devices, sexing systems and protocols for field evaluation for both *Anopheles* and *Aedes* species^30,31^. IIT relies on mass rearing and mass release of males depleted of their natural *Wolbachia* endosymbiont and harbouring a *Wolbachia* strain different from that present in the wild target mosquito population. *Wolbachia* is a gram-negative bacterium, a common symbiont of insects, including mosquitoes^32^, capable of inducing cytoplasmic incompatibility, i.e., mating between *Wolbachia*-infected males and wild females, without or with a different *Wolbachia* strain, results in embryonic lethality^33^. Pilot studies carried out in Kentucky (USA) and Italy support IIT as a valuable approach to suppress *Ae. albopictus* population^34,35^. Combining SIT and IIT and releasing millions of factory-reared *Ae. albopictus* adults over a two-year period on two small islands in Chinese city of Guangzhou, Zheng and colleagues have recently demonstrated that the near elimination of field populations of a mosquito specie is achievable^36^.

Nowadays, more than thirty SIT or SIT/IIT pilot trials are ongoing worldwide, mainly against *Aedes* mosquitos, and, very recently, IAEA and WHO released a guidance framework for the assessment of the feasibility of SIT as a mosquito-control tool against *Aedes*-borne diseases^37^ and a phased conditional approach guideline for mosquito management using SIT^38^. According to these guidelines, the testing and the implementation of Integrated Pest Management (IPM) techniques for mosquito control, including a SIT and/or IIT components, require, as essential premises, i) the selection of a proper and, possibly geographically isolated, site, ii) the baseline data on the bionomics of the infesting mosquito populations and iii) the local community engagements. In particular, a strong support and commitment of the local community and the early engagement of stakeholders, including residents in study sites and policy makers, are considered essential for the success of field trials^37^.

In this paper, we report on the characterization of the island of Procida in the Gulf of Naples (Italy), a Mediterranean site with a unique combination of instrumental features for the successful testing of innovative IPM approaches against *Ae. albopictus*. First the island has very suitable ecological and demographic features, i.e., i) a very small size (only 4.1 km^2^), ii) a completely urbanized and accessible territory, iii) high abundance of water containers in private gardens; iv) high human population density (2.83 inhabitants/ha - ISTAT 31/12/2018), and v) a significant infestation by *Ae. albopictus* initiated, according to Procida residents’ perception, around year 2000 (most probably introduced by tourists and/or maritime transport of goods) and grown into a serious nuisance problem in the last ten years. Second, many Procida citizens are familiar with SIT and of its advantages and effectiveness in insect pest control programs thanks to field performance tests of sterilized males of the Mediterranean fruit fly *Ceratitis capitata*, performed in the island during 1970’s and 1980’s, in a cooperative programme between the Italian National Committee for Research and Development of Nuclear Energy (ENEA) and the IAEA^39,40^. About 20 million sterile Mediterranean fruit fly males were released on the island from April to July 1986 leading to population suppression and protection of citrus fruits and therefore to a very positive perception of SIT by the residents thereafter.

Here, we present: i) baseline entomological data on the temporal dynamics and the spatial distribution of *Ae. albopictus* in the island collected through the active involvement of local citizens and administrators; ii) estimates of *Ae. albopictus* wild population size; and iii) estimates of dispersal and survival rate of irradiated *Ae. albopictus* males. These data provide insights on the bionomics of the Asian tiger mosquito in Southern Europe and draw attention to Procida Island as an ideal site for the planning of future pilot testing of innovative control programs against *Ae. albopictus* needed to foster future implementation in other Mediterranean and temperate areas.

## Results

### Temporal analysis

As a first step in the characterization of the *Ae. albopictus* population dynamics on Procida Island, we predicted the species oviposition pattern based on meteorological variables. A total of 44,244 *Ae. albopictus* eggs was collected by 26 ovitraps distributed all over the territory of Procida and Vivara islands (Supplementary Fig. S1 and Table S1 online) and monitored weekly from April 2016 to December 2016. The application of a generalized additive mixed model (GAMM) to assess the relationship between egg abundance and meteorological variables shows evidence of a strong effect of temperature on the number of weekly collected eggs (Fig. 1, Table1). On the other hand, the effects of cumulated rain and of average wind speed result low or not significant, respectively (Fig. 1, Table 1).

**Table 1.**
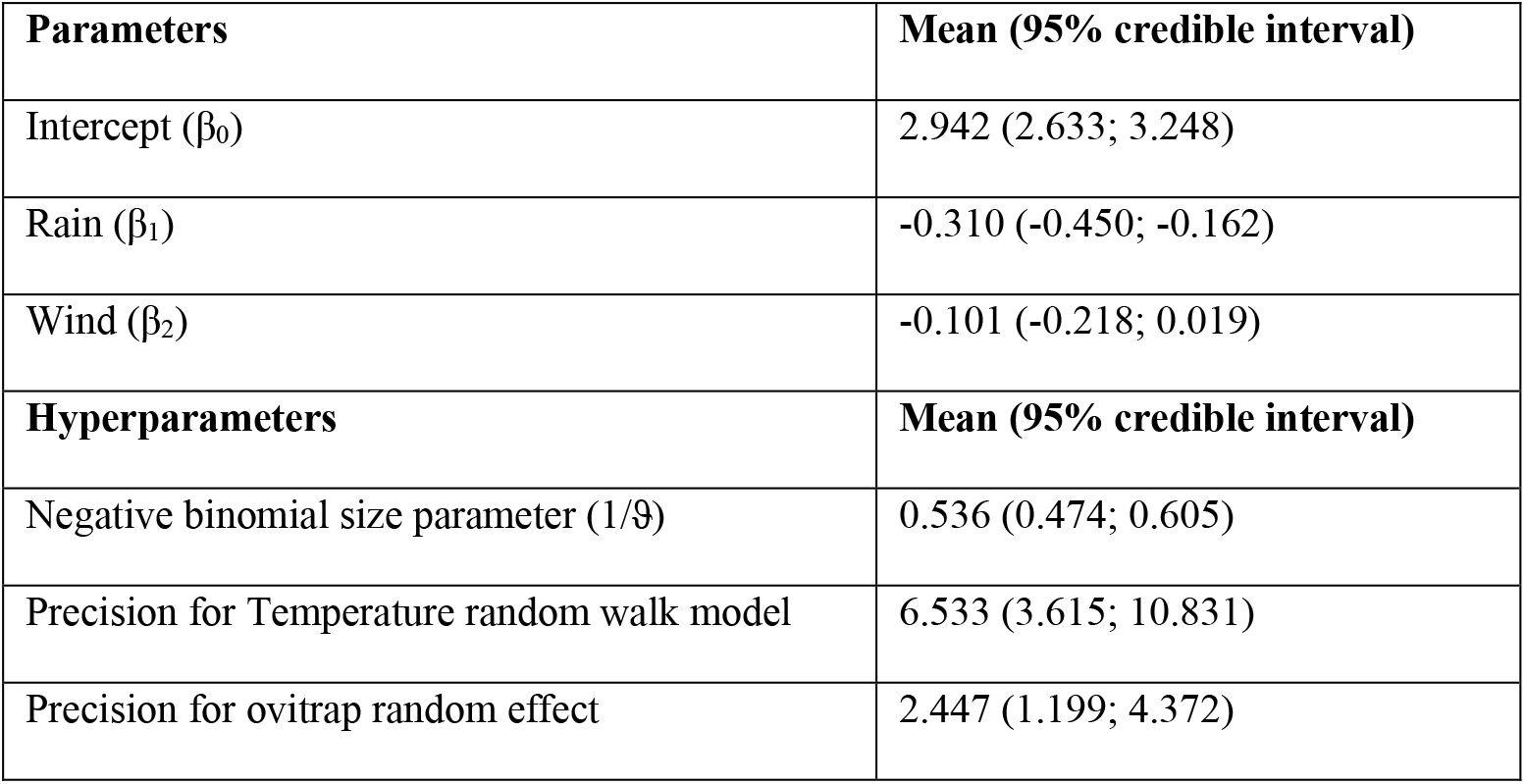
Results for the GAMM. Dependent variable is the count of eggs in ovitrap, independent variables are wind, rain considered as fixed effect and temperature as spline function, while the position of ovitraps as randomised effect. The posterior mean values and 95% credible intervals for both parameters and hyperparameters are provided. When the 95% credible interval includes zero there is no statistical support of a correlation between the independent and the dependent variable.

**Figure 1.**
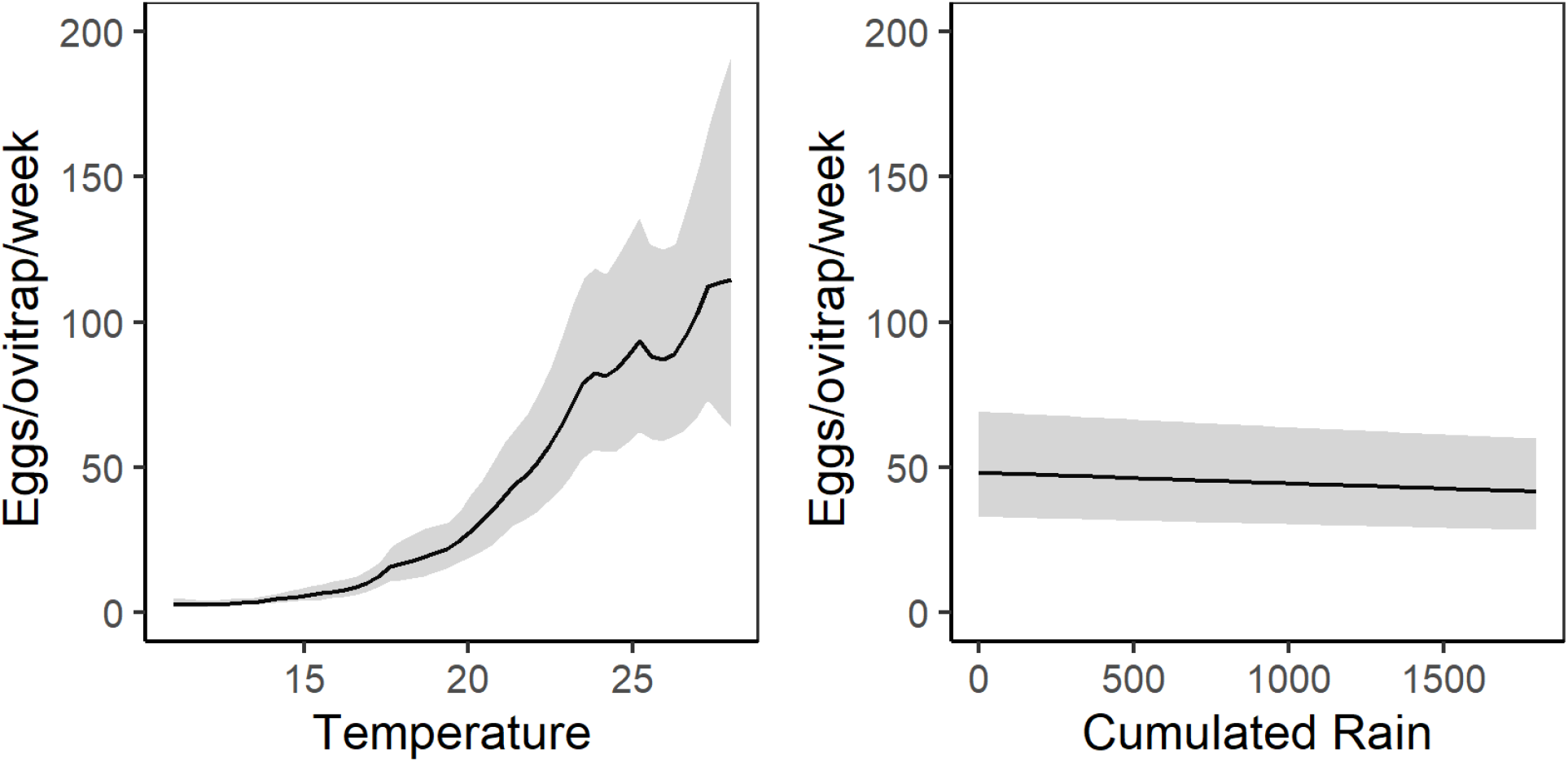
GAMM posterior predictive values of *Ae. albopictus* eggs/ovitrap/week in Procida Island. Left panel = temperature dependent mean value of eggs/ovitrap/week. Right panel = rain dependent mean value of eggs/ovitrap/week. Solid lines = GAMM posterior mean value of eggs/ovitrap/week. Grey areas = 95% credible interval. y-axis = number of eggs/ovitrap/week. x-axis (left panel) = weekly averaged temperature. x-axis (right panel) = weekly accumulated precipitation.

The posterior mean values and 95% credible intervals for GAMM parameters and hyperparameters are shown in Table 1.

Model assessment shows no-specific violation of the statistical assumptions (homogeneity, independence, autocorrelation, spatial correlation) (Supplementary Fig. S2 online). However, GAMM shows some under-dispersion (dispersion statistic = 0.71) under the assumption of a Negative Binomial distribution and a strong over-dispersion under a Poisson distribution (dispersion statistic = 39.3). Moreover, the model predicts a lower number of zeros compared to the observed (Supplementary Fig. S3 online). Model fit (GAMM model) estimated by Conditional Predictive Ordinate (hereafter CPO, see Materials and Methods, section Temporal analysis) and Bayesian p-value is poor (Supplementary Fig. S3 online), probably due to the large variation detected in ovitrap capture (range: 0-1500) coupled with a considerable proportion of zero captures (~25.7%). However, fitted values positively correlate with observed values (Pearson’s correlation: 0.697, df = 660, p-value <0.001) and the predicted oviposition temporal pattern is overall consistent with the observed one (Fig. 2).

**Figure 2.**
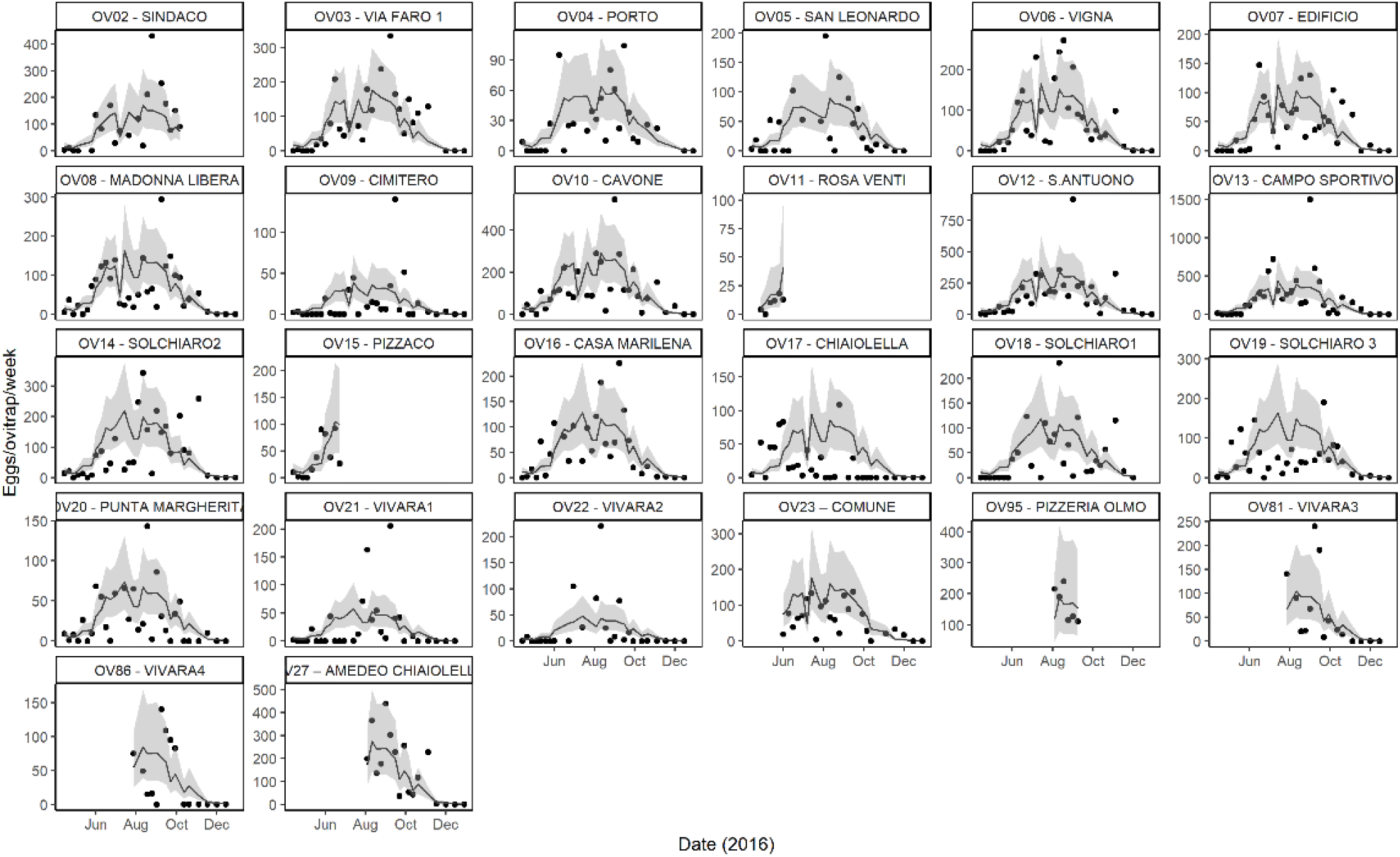
Observed and expected values *Ae. albopictus* eggs/ovitrap/week in Procida Island in 2016 estimated by GAMM model. Each panel represents a single ovitrap. Black dots = observed values of eggs/ovitrap/week; solid line = GAMM posterior mean value of eggs/ovitrap/week; grey area = 95% credible interval; x-axis = months of collections in 2016; y-axis = eggs/ovitrap/week; the scale differs per panel to help visualization.

Most relevantly for future implementation of innovative IPM in the island, GAMM simulations based on the mean number of eggs/ovitrap/week and meteorological data allow to estimate the start (e.g., 21^st^ April and 5^th^ May) and the end (e.g., 20^th^ October and 13^th^ October) of the breeding seasons in 2016 and in 2017, respectively (Fig. 3).

**Figure 3.**
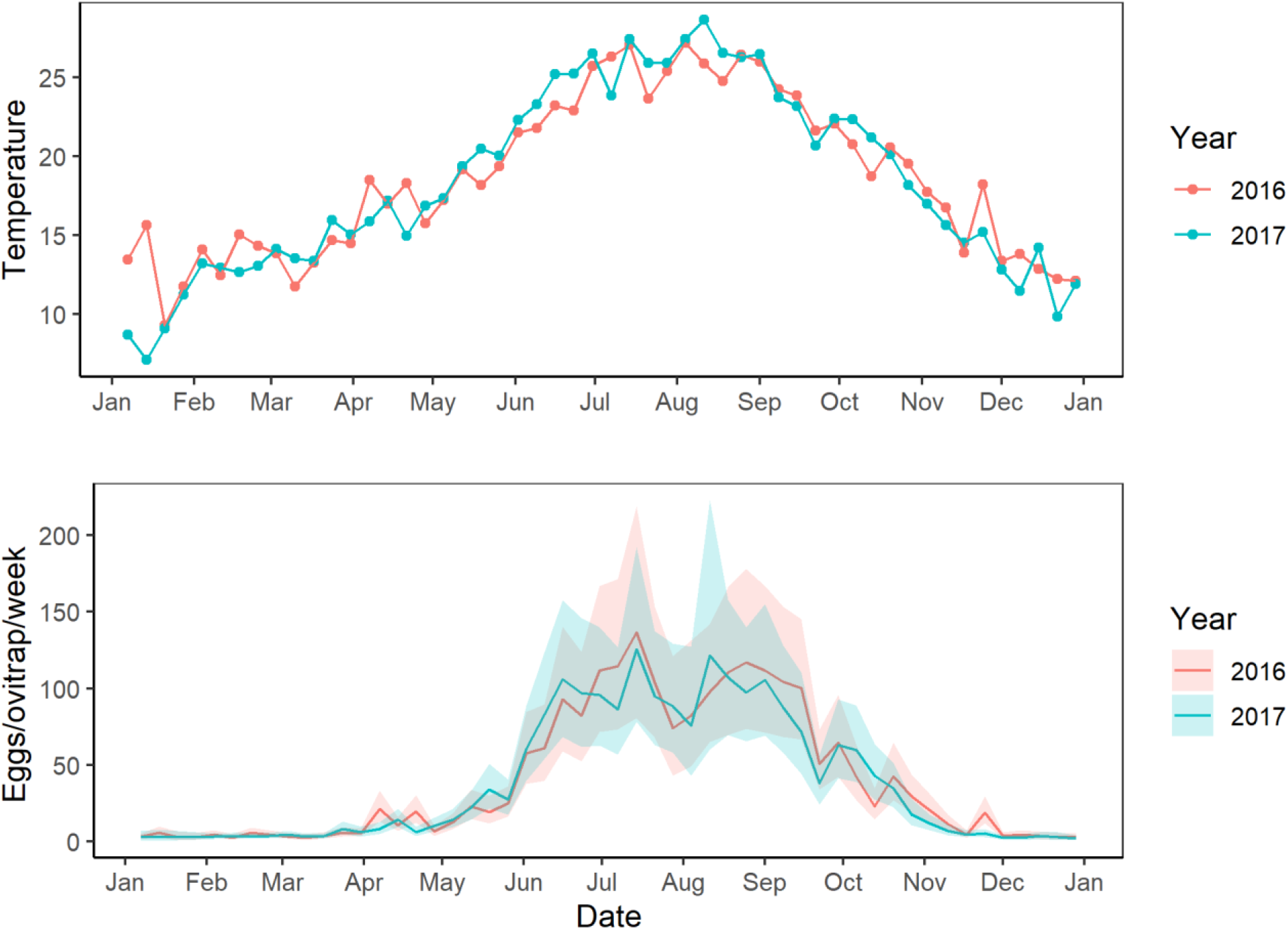
Temporal pattern of temperature (top) and *Ae. albopictus* eggs/ovitrap/week (bottom) in Procida Island in 2016 and 2017. Upper panel = average weekly temperature. Lower panel = eggs/ovitrap/week observed in 2016 and predicted in 2017. Solid lines = GAMM posterior predictive mean value of eggs in ovitrap, Shaded areas = 95% credible intervals.

### Spatial analysis

We used a geostatistical approach to analyse the spatial distribution of *Ae. albopictus* in Procida Island during the two predicted peaks of density of July and September, based on data from 101 ovitraps distributed across Procida Island and monitored in weeks 29-30 in July 2016 and 37-38 in September 2016. Geostatistical approaches, with particular emphasis on ordinary kriging, have been previously applied to study spatial distribution of insect vectors, e.g., to study either the association between vector distribution and pathogen transmission^41^ and the critical low temperature for the survival of *Aedes aegypti* in Taiwan^42^, or to relate entomological indicators to the incidence of dengue^43^ and to analyse seasonal and spatial distribution of *Ae. aegypti* and *Ae. albopictus* in San Paulo municipality (Brasil)^44^.

A total of 40,811 *Ae. albopictus* eggs were collected, 15,456 in weeks 29-30 and 25,355 in weeks 37-38. The first step in our geostatistical analysis was to carry out the variographic analysis to recognize the possible presence of a spatial structure in the data sampled, which in our case is the number of eggs/ovitraps/week (Fig. 4). The presence and type of spatial dependence was evaluated by means of the experimental variogram, which represents the pattern and the degree of spatial dependence between the sampling locations. The experimental variograms, obtained after a preliminary analysis of the cloud variograms (Supplementary Fig. S4 online), show a good spatial structure in the total number of eggs/ovitraps/week with a relatively high sill-nugget ratio (Supplementary Fig. S5 online). This underlines that an important part of the sample variance can be explained in terms of the spatial covariance and can be used, by means of kriging weights, to make spatial interpolation over a regular grid to get a continuous map of eggs deposited in the ovitraps.

**Figure 4.**
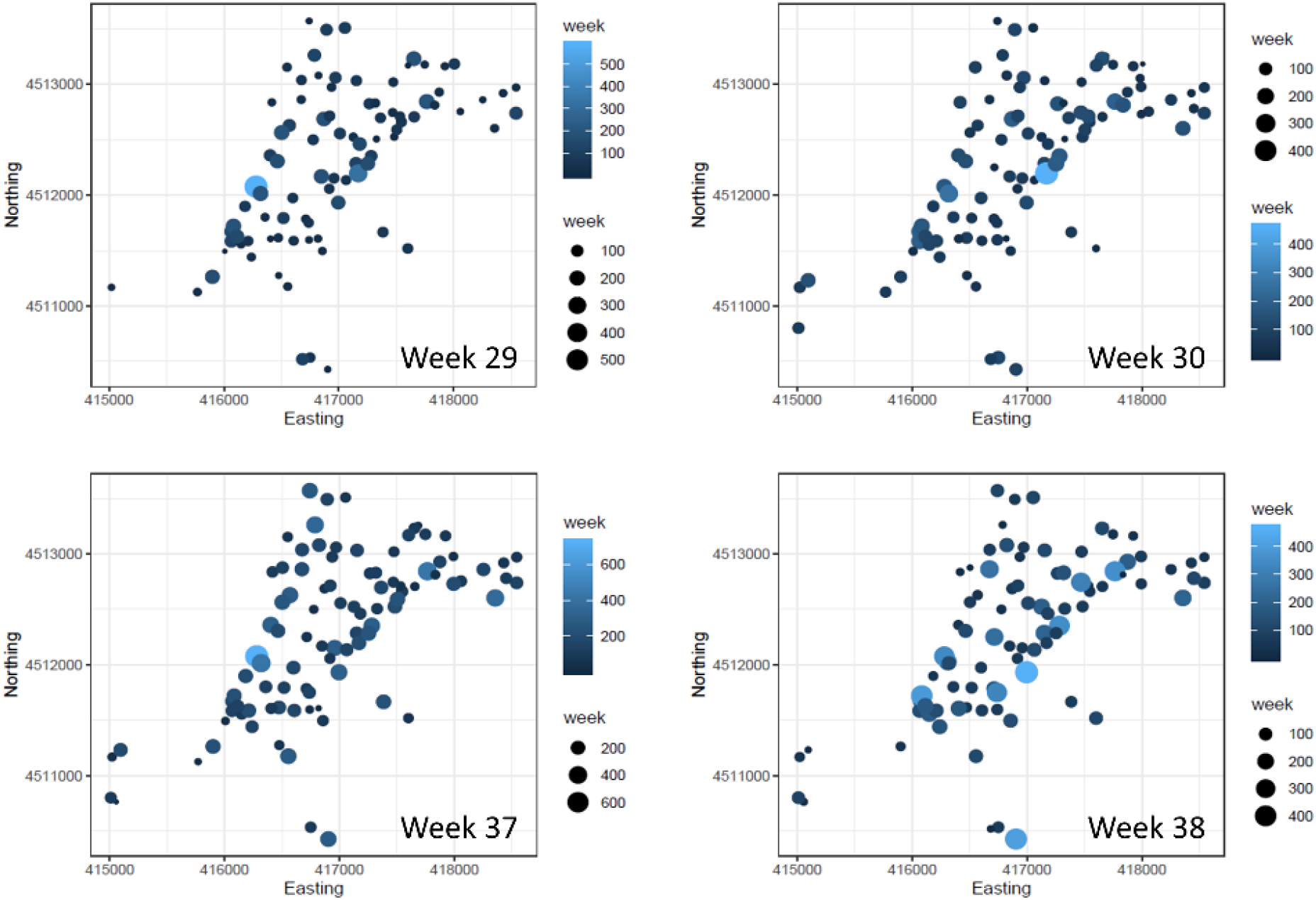
Number of *Ae. albopictus* eggs/ovitrap/week in Procida Island in July And September 2016. Spatial coordinates Northing and Easting are expressed in meters using the coordinate reference system with EPSG code 32633 (https://epsg.io/32633). Circle size and colour scales are proportional to the magnitude of collected eggs/ovitrap.

The ordinary kriging maps highlights a general uniform and wide presence of *Ae. albopictus* in the island, with some “hot spots” of oviposition activity (Fig. 5). We calculated a set of geospatial covariates that could aid the spatial analysis of target measurements (ovipositions) in a multivariate context. Different data sources were used, such as a digital elevation model, a map of land use and land cover and a map of distance from the sea. A geo-morphometric analysis of a digital elevation model was performed to calculate slope, aspect, topographic position index, terrain ruggedness index and roughness amongst others. Four rasters of growing degree days were calculated, one for each week, as a proxy of the spatial distribution of *Ae. albopictus* ovipositing females in Procida. All these covariates poorly contribute to the analysis of spatial variance of ovideposition, as highlighted by very low Pearson correlation values, ranging from −0.19 to 0.28, and no significant correlations are observed (Supplementary Tab. S2 online). This finding reinforces the conclusion of a high spatial uniformity of the Procida Island.

**Figure 5.**
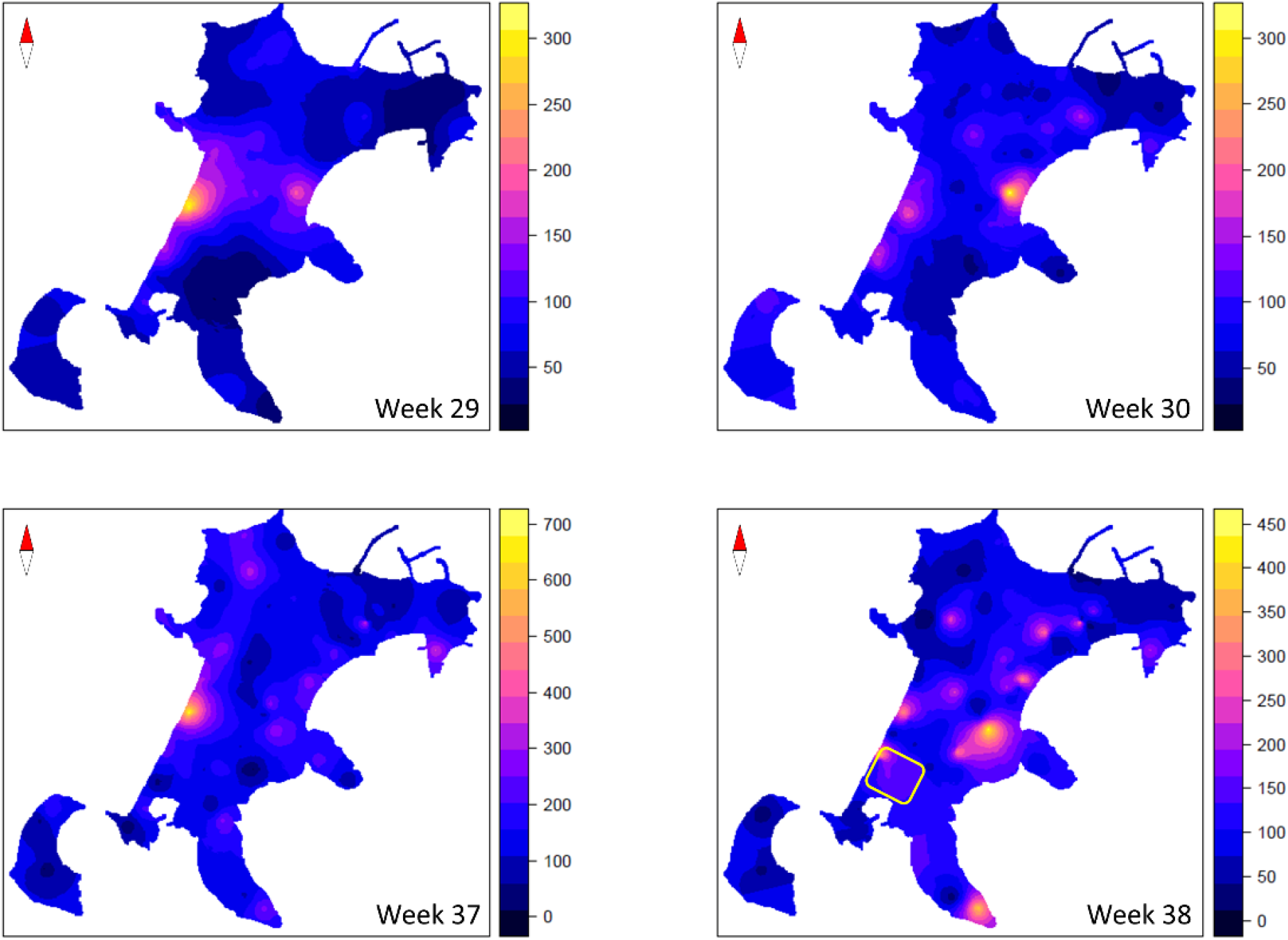
Ordinary Kriging maps of *A. albopictus* oviposition on Procida and Vivara islands. The figure shows the estimated ordinary kriging of *Ae. albopictus* oviposition on Procida Island in 4 weeks from July to September 2016. The colour gradient corresponds to the variation range of the estimated egg numbers. The yellow rectangle represents the area selected for the MRR experiments.

#### Ovitrap monitoring of *Ae. albopictus* on Vivara reserve

The data collected for the temporal analysis in Vivara island by four ovitraps monitored in April-December 2016 indicate the presence of *Ae. albopictus* also in this inhabited natural reserve, where potential hosts are represented by rodents, birds and reptiles. To better analyse the mosquito spatial distribution in the area, we further monitored, in collaboration with a Procida resident, expert guide of the Vivara reserve, 31 ovitraps located along the two main paths in the island during two high density periods in September 2018 (week 38) and July 2019 (week 28) (Supplementary Table S1 and Fig. S6 online). All ovitraps were found positive for *Ae. albopictus* eggs: a total of 18,891 and 6,608 eggs were captured in September 2018 (611 eggs/trap/week), and in July 2019 (213 eggs/trap/week), respectively. Adult collection by BG-sentinel traps baited with BG-lure yielded 7 males and 31 females by one BG trap/week in September 2018 and 19 males and 27 females by two BG-trap/week in July 2019, reinforcing ovitrap data showing high abundance of *Ae. albopictus* in Vivara natural reserve.

#### Mark-release-recapture experiments

We exploited results of Mark-Release-Recapture (MRR) experiments to estimate the density per hectare of local *Ae. albopictus* males during the high-density peak in September 2018 and to evaluate the dispersal capacity of sterile males. Thirty-nine sampling stations were located in four concentric annuli around the release site in the touristic district of “La Chiaiolella” (~3.1 stations/hectare) in the southern part of the island (Fig. 5 and Fig. 6).

**Figure 6.**
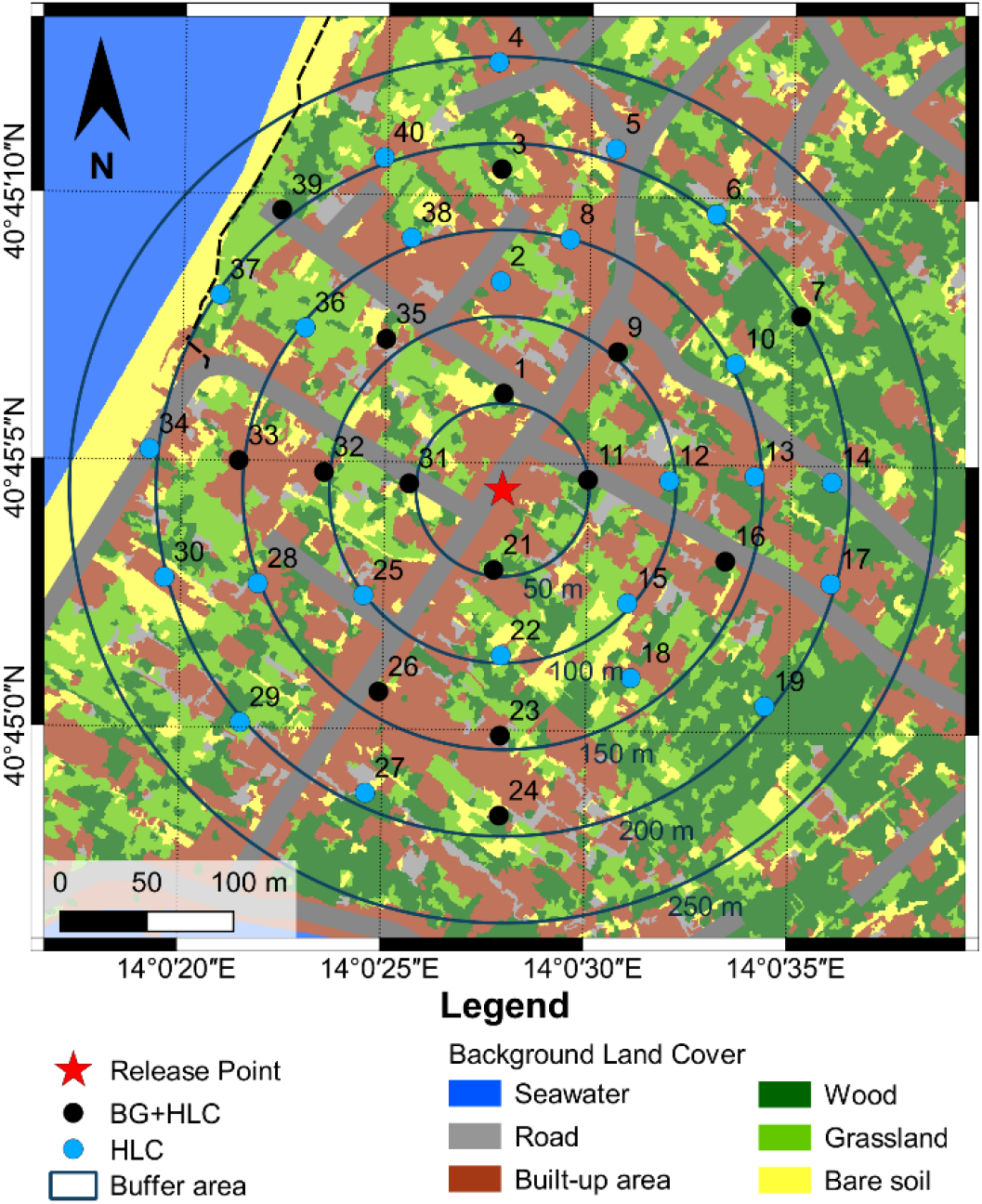
Distribution of *Aedes albopictus* sampling stations in the mark-release-recapture study site in “La Chiaiolella” (Procida). Red star = release site. Black circles = HLC + BG-traps recapture stations. Light blue circles = HLC recapture stations. Circles = 50 m-annuli around release site (Google maps).

In total, 7,836 (MRR1) and 9,680 (MRR2) marked sterile *Ae. albopictus* males (PRO1 strain) were released in September 2018. In MRR1, 2,009 wild males and 169 marked males were recaptured by HLC (N marked males=79) and by BG (N=90) during 14 days of collection, for a recapture rate of 1.8% (N=143) in the first 6 days. Among these, 97.8% were recaptured within 50 m from the release site and none at ≥150 m from it (Table 2). In MRR2, 2,319 wild males and 165 marked males were recaptured by HLC (N=111) and BG (N=54) during 6 days of collection, for a total recapture rate of 1.7%. Among marked males, 87.9% and 6.7% were recaptured within 50 m and at ≥150 m distance from the release site, respectively (Table 2). Overall, only 4% and 9% of recaptured males were collected by BG sentinel and HLC beyond 50 m from the release point, respectively (Table 2).

**Table 2.**
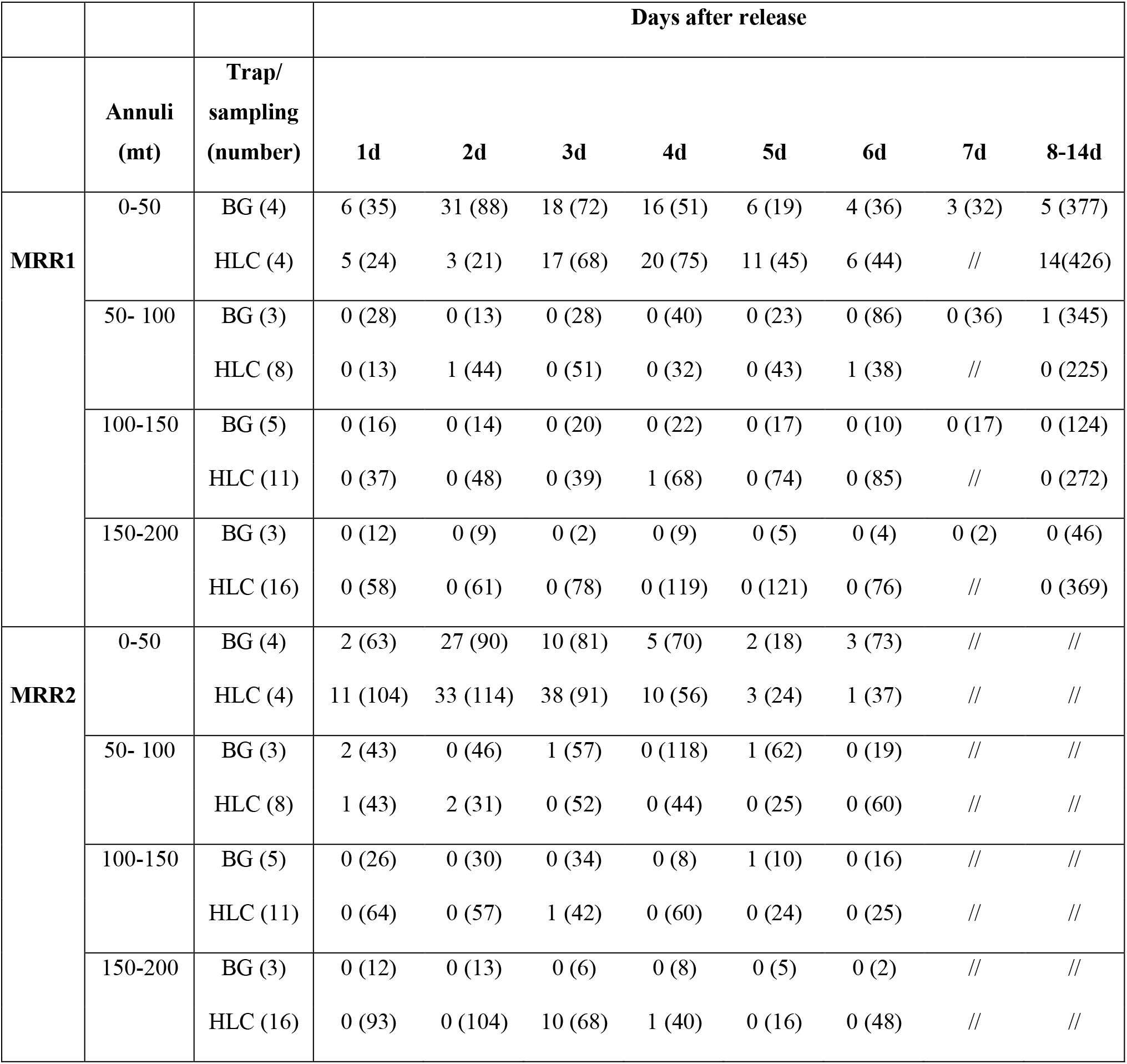
Number of marked (and unmarked) *Ae. albopictus* males recaptured in two Mark-Release-Recapture (MRR) experiments in Procida Island. BG = BG-sentinel traps (number of BG within each annulus); HLC = Human Landing Catches (number of HLC sites within each annulus).

Estimates of cumulative MDTs (1-6 days) was 51 m in MRR1 and 61 m in MRR2 based on BG-trap collections, and 52 m in MRR1 and 57 m in MRR2 based on HLCs (Table 3).

**Table 3.** Mean distance travelled of recaptured *Aedes albopictus* sterile males in two mark-release-recapture experiments in Procida Island.

|  | MRR1<br>(BG) | MRR2<br>(BG) | MRR1<br>(HLC) | MRR2<br>(HLC) |
| --- | --- | --- | --- | --- |
| day after<br>release |  |  |  |  |
| 1 | 51 | 94 | 51 | 55 |
| 2 | 51 | 51 | 63 | 57 |
| 3 | 51 | 61 | 51 | 61 |
| 4 | 51 | 51 | 53 | 55 |
| 5 | 51 | 89 | 51 | 51 |
| 6 | 51 | 51 | 58 | 51 |
| 1/6 days | 51 | 61 | 52 | 57 |

Population size was estimated over areas of either 50 m (estimated flight range) or 200 m (whole sampling area) radius (Table 4 and Fig. 7): results predict that the number of males/ha is higher within 50 m-radius area than in the whole one. This effect is probably due to heterogeneities in mosquito abundance (as estimated by field collection), which increases slower than the size of the area (which increases quadratically). Estimates of mosquitoes/ha based on recaptures in the 50 m-radius area largely differ depending between GLM and Fisher Ford methods (Table 4 and Fig. 7). The estimated survival parameter of marked mosquitoes ranges from 0.8 to 0.95 according to the sampling area and the trap type considered in the estimation.

**Table 4:** Wild *Aedes albopictus* population size in Procida Island and survival of sterile males estimated based on Mark Release Recapture (MRR) experiments within either a 50m- and a 200m-areas around release site. Density/ha = number of males/ha; N = estimate of population size/day; λ = survival rate estimated by logistic regression; confidence intervals based on 1000 bootstrap replicates with the method of percentile bootstrap at 95% level; m = number of marked mosquitoes recaptured; n = total number of mosquitoes captured (marked+wild).

| MRR | Methods | Type of traps | Annuli (m) | N | Density per ha | n | m | $\lambda$ | 95% CI per density per ha |
| --- | --- | --- | --- | --- | --- | --- | --- | --- | --- |
| 1 | Regression Model | BG | 50 | 24692 | 31437 | 301 | 81 | 0.92 | 20753-48490 |
|  |  |  | 200 | 39361 | 3132 | 659 | 81 | 0.8 | 2173-4618 |
|  |  | HLC | 50 | 32013 | 40761 | 277 | 62 | 0.87 | 22786-76221 |
|  |  |  | 200 | 144447 | 11494 | 1350 | 65 | 0.95 | 7075-19673 |
|  | Fisher-Ford(a) | BG | 50 | 14727 | 18750 | 301 | 81 | \\| | 13048-24835 |
|  |  |  | 200 | 11659 | 928 | 659 | 81 | \\| | 485-963 |
|  |  | HLC | 50 | 27058 | 34450 | 277 | 62 | \\| | 29223-47132 |
|  |  |  | 200 | 122611 | 9757 | 1350 | 65 | \\| | 9301-10989 |
| 2 | Regression Model | BG | 50 | 53928 | 68663 | 395 | 48 | 0.81 | 43566-112028 |
|  |  |  | 200 | 110256 | 8774 | 910 | 53 | 0.8 | 5753-13914 |
|  |  | HLC | 50 | 39481 | 50267 | 426 | 96 | 0.94 | 36021-71275 |
|  |  |  | 200 | 88389 | 7034 | 1322 | 114 | 0.86 | 5333-9425 |
|  | Fisher-Ford(a) | BG | 50 | 16618 | 21158 | 395 | 48 | \\| | 16810-31257 |
|  |  |  | 200 | 339776 | 3165 | 910 | 53 | \\| | 2700-4276 |
|  |  | HLC | 50 | 27024 | 34408 | 426 | 96 | \\| | 21814-41779 |
|  |  |  | 200 | 42421 | 3375 | 1322 | 114 | \\| | 2526-3367 |

**Figure 7.**
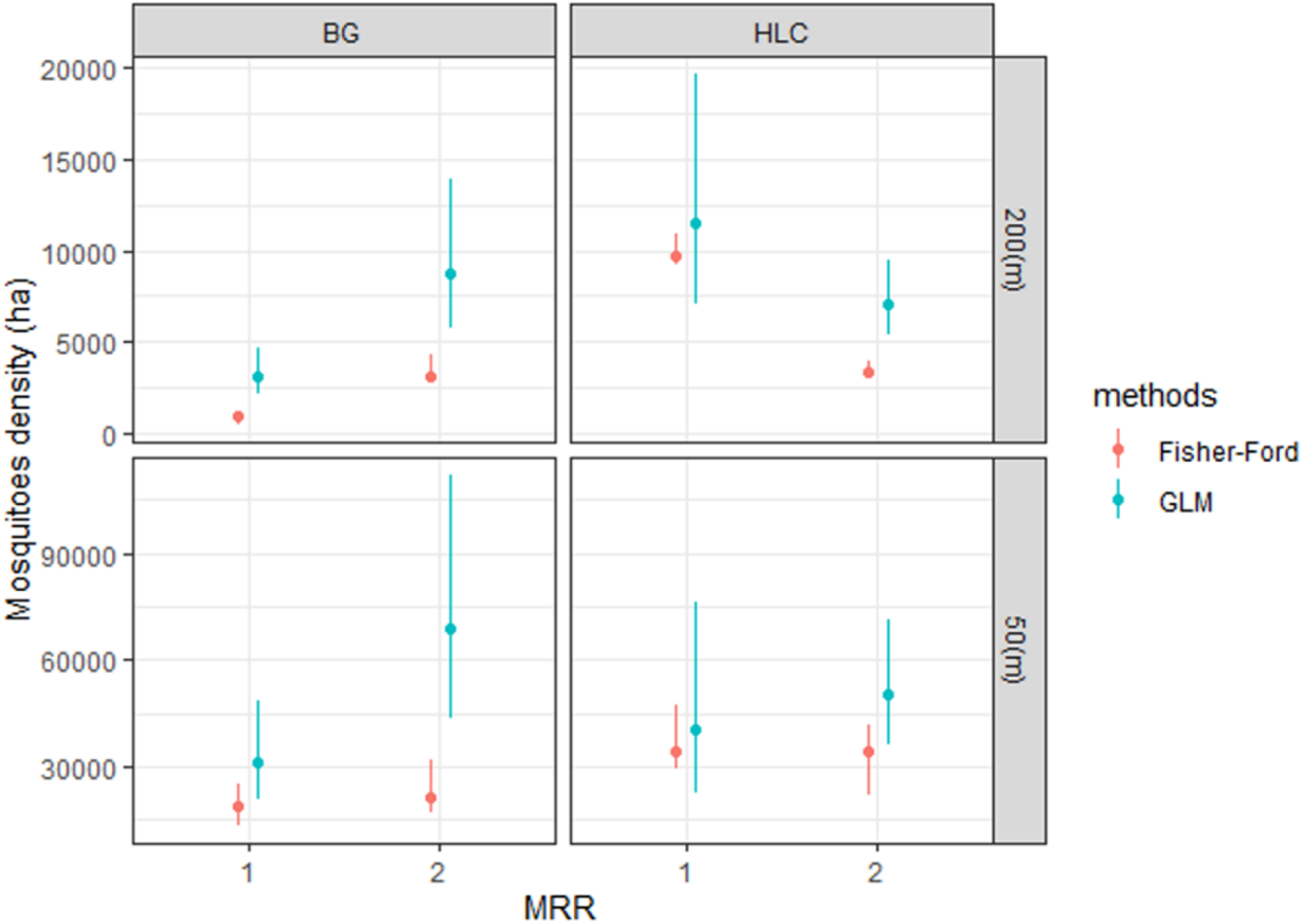
Wild *Aedes albopictus* population size in Procida Island estimated based on Mark Release Recapture (MRR) experiments within either a 200m- and a 50m-areas around release site. Blue lines = GLM-based estimates; red lines = Fisher-Ford based estimates; dots = mean value of mosquitoes/hectare; vertical segments = 95% confidence intervals using percentile bootstrap (red), using equation [5, supplementary Methods S1 online] (blue).

#### Community-engagement assessment

In September 2015, as a first step in our community engagement strategy, we conducted a public survey among Procida residents on their knowledge on *Ae. albopictus*-associated public health risks, on their actions to reduce mosquito nuisance and reproduction and on their interest to support and to participate to monitor and control campaigns (Table 5). Among the 200 randomly selected people interviewed (corresponding to about 2% of the resident population), 77% resulted aware of the capacity of mosquitoes to transmit diseases, 85% declared to use electric diffusers, mosquito nets or chemical repellents to protect themselves from bites, 11% removed water containers from their properties and only 3% use larvicide products in standing water. The large majority of interviewed people was in favour of a mosquito control programme in Procida, but a minority of them agreed to contribute economically (33%) or to participate as volunteers in the control programme (25%).

**Table 5.** Results of surveys carried out among Procida Island residents in 2015 (N=200) and 2019 (N=191).

| Survey question | Responses 2015 (%) |  | Responses 2019 (%) |  |
| --- | --- | --- | --- | --- |
|  | YES | NO | YES | NO |
| Do you know that the Asian tiger mosquito can transmit viral diseases to humans? | 77 | 23 | 73 | 27 |
| Do you use protective measures against mosquitoes? | 85 | 15 | 91 | 9 |
| Do you use electric diffusers? | 58 | 42 | 37 | 63 |
| Do you use mosquito nets? | 54 | 46 | 71 | 29 |
| Do you use insect repellents? | 45 | 55 | 47 | 53 |
| Do you use larvicides? | 3 | 97 | 2 | 98 |
| Do you remove standing water? | 11 | 89 | 11 | 89 |
| Would you welcome a regional/municipal mosquito control programme? | 88 | 12 | 96 | 4 |
| Would you agree to the installation in your property of traps for the capture and monitoring of mosquitoes? | 44 | 56 | 78 | 22 |
| Would you agree to contribute personally to the financing of a mosquito control project? | 33 | 67 | 54 | 46 |
| Are you interested in participating, as a volunteer, to a mosquito monitoring and control programme in Procida? | 25 | 75 | 33 | 67 |

In September 2019, we administered again the same questionnaire to 191 randomly selected residents to record possible changes in people feedback after our research activities. While no significant changes were observed with reference to people knowledge in the mosquito and preventive/protective measures taken (with the only exception of an apparent increase in the use of mosquito nets), we observed an increased interest in mosquito control program and an increased availability in supporting or participating to monitoring and control actions in the island.

## Discussion

The results here obtained provide estimates of baseline data on *Ae. albopictus* population in Procida (Italy) which could facilitate possible experiments in the island aimed to assess the feasibility and potential effectiveness of SIT/IIT-based control methods in Mediterranean regions and testify the possibility and the value of creating synergies between research groups, local administrators and citizen. The data obtained also add to the present knowledge on the bionomics of wild *Ae. albopictus* in the Mediterranean region and of irradiated males to be exploited for SIT.

First, *Ae. albopictus* seasonal activity in Procida is shown to span from late April/early May to October/early November, as already reported from Rome and Lazio region^45,46^, while data from northern Italian regions indicate a <6 months-long reproductive season^47^. Based on the estimate values obtained by GAMM the egg abundance dynamics in Procida is shown to be dependent on temperatures (Table 1; Fig 1) and is characterized by two peaks, similarly to the temporal pattern reported from Rome^46^. It is important to note that the results obtained by the temporal analysis had a limitation due to monitor the dynamic of population for just one year and have been estimates by ovitrap capture known to have large variation in collections. Based on these evidences SIT and/or IIT-based interventions in Procida should start not later than April to have their maximum efficacy. Moreover, these baseline data on *Ae. albopictus* seasonal dynamic in the island could be used to assess the effectiveness of SIT and/or IIT control activities, by comparing not only the absolute mosquito densities but also their temporal trend in the presence and in the absence of these interventions. If validated with further field data, the GAMM model utilized in the present paper could represent a trustable predictive model exploiting locally collected meteorological data to define more precisely the starting period of control interventions.

Second, the analysis of the spatial distribution of the *Ae. albopictus* highlights an overall uniform and massive presence of the mosquito in the island, in agreement with the perception of the researchers that collected the data during the monitoring campaign. This is consistent with the nature of the island territory, completely urbanized and fragmented in hundreds of private properties with residential buildings surrounded by gardens with vegetable cultivations and/or orchards with citrus plants and family-type farming of animals. Indeed, it was very easy to identify several anthropic water sources ideal for the species larval breeding in all sites selected for ovitrap positioning. Interestingly, high *Ae. albopictus* densities are also observed in the inhabited natural reserve of Vivara, indicating the mosquito population of Procida maintains the capacity to colonize sylvatic environments. This observation highlights the importance to also consider these kinds of habitats for the proper planning of future mosquito control interventions in Procida, as well as in other Mediterranean areas.

Third, we estimated the mosquito population size at the end of the reproductive season (September) based on results of MRR experiments, to provide a numerical value to the citizen’s and researchers’ perception of very high densities/nuisance. Estimates based on 200m-dispersal (928-9,757 males/ha with Fisher-Ford and 3,132-11,494 males/ha with regression) are comparable with the values estimated in the north of Réunion island (i.e., 639 males/ha in September 2016 to 6,000 males/ha in December 2015)^48^ and higher than estimates based on MRR experiments conducted in Rome in September 2009 (i.e. 135 mosquito/ha with regression and 294 m/ha with Fisher-Ford)^49^. Higher population size values (18,750-68,663 male/ha) are estimated by both statistical methods applied (despite GLM provided greater values than Fisher-Ford equation), under the assumption of a 50m dispersal. Although, population size estimates are to be taken with caution as they are affected by assumptions made (e.g. on dispersal), as well as by methodological approaches (e.g. releases of males or females, recapture method, statistical method), the very high number of *Ae. albopictus* individuals estimated in Procida underlines the capability of this species to reach very high population densities with detrimental effect on the quality of life of residents and tourists and, hence, the overall tourism-centred economy of the island.

Fourth, results clearly show the success of the effort of the research groups in working in synergy with public administration and citizen in Procida. A detailed community engagement strategy was developed (see Methods and Supplementary Figure S7 online), including: i) preliminary engagement of the local administrators; ii) selection of a first group of twelve volunteers interested to participate in *Ae. albopictus* monitoring by personally dealing with ovitrap management over one year; iii) active participation of many citizen to allow and easy capillary access to the whole island territory. This allowed optimization of the monitoring and experimental procedures and the possibility to carry out the activities with a limited budget. Indeed, according to the surveys carried out in 2015 and 2018, Procida citizen involvement in mosquito monitoring and control can be further increased. The limited commitment of the residents in standing water elimination and in larvicide usage highlights the need to increase knowledge on best practices to be implemented at the individual level to reduce mosquito reproduction (e.g., by specific educational project in schools on mosquito biology and control) as a relevant step to lower mosquito densities and creating a situation more suitable for the success of pilot SIT/IIT studies and public mosquito control activities.

Fifth, results of MRR experiments show a short-range dispersal of irradiated males, likely favoured by the uniform landscape in Procida Island with plenty of resting and swarming sites for adult males. However, the <60 m mean daily distance travelled by the males is in the range of values estimated for non-irradiated *Ae. albopictus* males in Switzerland^50^ and Reunion Island^48^ although lower than estimated for *Ae. albopictus* females in Italy^51,52^ and Switzerland^50^. This suggest that the flying capacity of release males is not affected by manipulations, irradiation and transportation, but also highlights the need of planning multiple short-distance releases within a SIT intervention schemes.

Sixth, daily survival estimates of irradiated males ranged between 0.80 and 0.95, which are in the range of estimates for not-irradiated males in Missouri (USA) (survival rate=0.77)^53^; Reunion Island (survival rate =0.90^48^ and 0.94-0.95^54^) and Switzerland =0.88^50^. This suggest that not only the dispersal capacity, but also the longevity of release males is not affected by manipulations, irradiation and transportation, supporting the potential effectiveness of SIT interventions based on male production and sterilization in centralized facilities, even if distant from intervention area.

## CONCLUSIONS

The results obtained significantly advance in the path toward the feasibility of a demonstrative project to assess the cost-effectiveness of mosquito control program integrating conventional control method (e.g., larval source removal and larvicide treatments) with SIT and/or IIT in the fight against *Ae. albopictus* and *Aedes*-borne diseases in Mediterranean Europe and in other temperate regions. The relevant nuisance and threat to tourism and health caused by the mosquito in the island and the positive feedback shown by the local stakeholders (i.e., administrators and citizen) in being active parts in innovative mosquito control projects represent major assets in the selection of the island as study site. The acquired knowledge of the species spatial distribution, seasonal dynamics and absolute abundance obtained by 4 year-research activities in the island provide the ground for evidence-based planning of where, when and how to implement the interventions and for assessment of their effectiveness. For instance, the homogeneous distribution of *Ae. albopictus* population in the urbanized part of the island as well as in the wildest part (i.e., the Vivara natural reserve) points to the need of an island-wide capillary release of males, which could be achieved by drones ^55^ and/or by a large network of volunteers involved in ground releases. Indeed, the success of a such pilot experiment, performed in a small island with urban-like human density, could be and excellent endorsement and proof-of-principle for the implementation of SIT and/or IIT against *Aedes* mosquito in wider Mediterranean areas.

## Methods

### Study site

The island of Procida belongs to the Phlegraean archipelago and it is situated in the Naples gulf (Italy). It is a 3.7 km^2^ large flat volcanic island (average 27 m above sea level) with a 16 km-long rough coastline which forms four capes. A 0.4 km^2^ large satellite island, Vivara, a natural uninhabited reserve, completes the Procida territory. Procida is a highly urbanized with a completely accessible territory and a very high population density of 2.83 inhabitants/ha (ISTAT 31/12/2018); population density approximately doubles during summer months, because of tourism, which represents the main local economic activity. Most of Procida’s private properties include a garden with ornamental flowers, vegetable cultivations and/or orchards with citrus plants and family-type farming of chickens and rabbits. The climate is temperate with an average annual temperature of 16.2 °C and an average annual precipitation of 797 mm (https://en.climate-data.org/) and it is classified as Csa by Köppen and Geiger (Csa = Hot-summer Mediterranean climate)^56^.

### Ovitrap collections for temporal and spatial population dynamics

Cylindrical black plastic jar, 15 cm high, 12 cm in diameter with an overflow hole at 8 cm from the base, were utilized as ovitraps. Ovitraps were filled with about 600 ml of tap water and heavy-weight seed germination paper strips (Anchor Paper Co. USA) were utilized as oviposition substrate, with strip of 30 x 9 cm, lined to the internal wall of the ovitrap. Ovitraps were located at ground level, in shaded areas near the vegetation. For the temporal analysis, 26 ovitraps were distributed all over the island and monitored weekly from 2016-04-14 to 2016-12-31 (Supplementary Fig. S1 and Table S1 online). For the spatial analysis, 75 additional ovitraps were deployed across the island and monitored, together with the previous 26, for two weeks in July 2016 (weeks 29-30) and two weeks in September 2016 (weeks 37-38). Collected germination paper strips were brought to the laboratory and eggs counted under a stereomicroscope.

### Citizen science and community engagement strategy

All experimental activities were performed in strict collaboration with local administrators and citizen in Procida, following the workflow described in Supplementary Figure S7 online. As a first step, with the help of local administrators we selected twelve volunteers interested to participate in an *Ae. albopictus* temporal dynamic monitoring using ovitraps (Supplementary Fig. S8 online). A one week-long workshop led by expert operators was organised on site to train the volunteers on how to manage ovitraps and collect germination paper strips to be weekly delivered to an expert technician for egg counting. As a second step, with the help of Procida administrators and of the twelve volunteers, 79 families were selected and allowed access to their private properties to place the ovitraps required for the spatial analysis, which were managed by expert operators for 4 weeks (Supplementary Fig. S8 online). In the third step, we involved the local community of “La Chiaiolella” area for the MRR experiments with the help of a “facilitator”, a well-respected man in the community recommended by Procida administrators. Thanks to his help, twenty families, local merchant associations and parishes agreed in hosting in their private properties the re-capture stations (see below) and two field laboratories for managing of the sampling instruments and the egg/adults counting (Supplementary Fig. S8 online). During the four years of the project (September 2015 – September 2018), several additional public activities were performed to further promote the public awareness and participation of the Procida community to our experiments (Supplementary Fig. S7 online), including the two public survey described in the results section.

### Production of sterile males

Experimental sterile *Ae. albopictus* male mosquitoes were produced from eggs collected in June 2018 by ovitraps within the study area. Eggs were transported at the laboratory of Sanitary Entomology and Zoology at the Centro Agricoltura Ambiente “Giorgio Nicoli” in Crevalcore (CAA - Bologna, Italy) and utilized to set a Procida *Ae. albopictus* strain named PRO1. PRO1 strain was reared as previously described^28^ in a climate-controlled insectary (28 ± 1°C, 80 ± 5% RH, 14:10 h L:D photoperiod). CAA Laboratories are certified with ISO9001, ISO14001 and ISO45001. Larvae obtained after standardized hatching procedures^27^ were reared at a fixed larval density (2 larvae/ml) and fed with a standard diet integrated with brewer yeast (IAEA-BY) liquid diet (5.0% w/v) at a mean daily dose of 0.5 mg/larvae^57^ for the first four days of development. Pupae were harvested once per cycle at about 24 h from the beginning of pupation and males were separated using a 1,400-micron sieve (Giuliani Tecnologie S.r.l., Via Centallo, 62, 10156 Torino, Italy). Male pupae were aged 24 h before being subjected to the irradiation treatments at the Medical Physics Department of St. Anna Hospital (Ferrara, Italy) using a gamma irradiator (IBL 437C, CIS Bio International, Bagnols sur Ceze, France; 65.564 TBq 1772 Ci ± 10% Cs-137 linear source) at a dose of 35 Gy and a dose rate of 2.1 Gy/min (± 3.5%). Following the irradiation procedure, no loss of pupae was observed. Irradiated pupae were then placed in petri dishes (12 cm diameter) filled with water and transferred in cardboard boxes (12 × 12 × h 18 cm), provided with additional separators to increase the resting areas, closed at the top with mosquito net, for emergence and shipment. Cotton pads soaked with 10% sugar solution were provided and secured at the top of each box to assure adult nourishment. Each box contained ca 1,500-2,000 adults and provided a vertical resting surface area of 1.3-1.0 sqcm per adult. Mosquito boxes were maintained at about 21 °C for the first two days after adult emergence and then transferred inside larger polystyrene container with adequate quantity of gel packs (phase change material) with the aim to maintain a temperature between 10 and 15 °C during shipment. About 10,000 adult sterile males were sent via ground transportation by express courier in two different expeditions in boxes of about 2,000 or 1,500 sterile males in the first and second release, respectively.

### Mark-release-recapture experiments

Sterile male releases were performed on 14 (MRR1) and 21 (MRR2) September 2018, at 15:00 PM (40°45’04.6’’N, 14°00’27.5’’E). Temperature and relative humidity at the release sites were 28°C and 59% and 27°C and 61% during MRR1 and MRR2, respectively. Immediately before release, males were marked with a coloured fluorescent powder (PROCHIMA s.r.l., Calcinelli di Saltara (PU), Italy) using manual insufflators to create a dust cloud inside the cardboard transportation boxes. A purple and green dye was used in MRR1 and MRR2, respectively, to differentiate males released in the two releases. Dusted 5 days-old sterile males were released by placing and opening the cardboard boxes in a sunny area without vegetation to favour dispersal. The cages were gently shaken for about 30 min, to induce the males to exit. The males that remained in the cage after 30 min were counted and deducted from the total. Recaptures began approximately 24 hours after each release and performed daily for 13 consecutive days in MRR1 and for 6 consecutive days in MRR2. Recaptures were performed in 39 sampling stations distributed in four concentric annuli around the release point and at 50 meter-distance from each other (Supplementary Table S1 online) selected in collaboration with the Procida administration and with 20 local families. At each sampling station, recaptures were performed by BG-Sentinel traps (operating continuously) and by Human Landing Catches (HLC) during the late afternoon peak of *Ae. albopictus* activity (indicatively from 4:30 to 7:30 PM). BG-Sentinel traps baited with BG-Lure were placed at ground level in shaded locations close to domestic areas. HLC were carried out by expert operators (co-authors of the present paper) using locally-made battery-powered hand-held electric aspirators for 15’ in at each station. Field-collected mosquitoes were identified using morphological keys and marked males were detected under stereomicroscope and UV lamp.

### Ethics declarations

This research was approved by the University of Naples Federico II CSV Ethics Board (Protocol # PG/2020/0090230) and by the Municipality of Procida with municipal resolution n°52 of 07 July 2016. All procedures performed in studies involving human participants were in accordance with the ethical standards of the institutional and/or national research committee and with the 1964 Helsinki declaration and its later amendments or comparable ethical standards. Informed consent was taken from all the participants involved in the study. All the people (co-authors of the present manuscript and Procida volunteers) present in Supplementary Fig. S8 online, gave their consent to the publication of the images. All authors declare no competing interests.

### Eco-climatic parameters

A spatial dataset to analyse distribution of descriptive land cover, geomorphological and climatic variables in Procida Island has been generated using open-source Geographic Information System, specifically GRASS GIS^58^ for data processing and spatial analysis and Quantum GIS^59^ for spatial analysis and layout generation.

Land cover variables were retrieved from supervised classification of digital multispectral aerial imagery collected by optical sensor in the visible spectrum on 16 June 2016 and 07 May 2011 at 0.5 m spatial resolution (Source: Italian National Geoportal, http://www.pcn.minambiente.it/GN/), using the methodology described in Manica et al. (2016). Mapped land cover classes were ‘trees’, ‘grasslands’, ‘roads/concrete’, ‘buildings’, ‘bare soil’, ‘water bodies’, ‘seawater’. Two main classes are derived from the land cover classified map: artificial surfaces’ (including ‘roads/concrete’ and ‘buildings’) and ‘natural cover’ (including ‘wood’, ‘grassland’, ‘bare soil’).

Topography of Procida Island is described in the Digital Terrain Model (DTM) at 2 m spatial resolution, generated from LiDAR acquisitions (Source: Italian National Geoportal, http://www.pcn.minambiente.it/GN/). The following additional geomorphological descriptors have been computed from DTM data using GDAL library (GDAL/OGR contributors, 2020): slope, aspect, terrain roughness, Topographic Position Index (TPI) and Terrain Ruggedness Index (TRI).

In the context of climatic variables, daily spatial maps at 30 m spatial resolution of air temperature climate variable in Procida has been computed combining *in-situ* measured meteorological data and satellite estimated Land Surface Temperature (LST). LST data represent the estimation of skin temperature detected at earth surface by remote sensing sensor. Meteorological data were downloaded from the ‘Ciraccio - INAPROCI2’ weather station using the Weather Underground database (https://www.wunderground.com/). Since daily map of LST estimates were not available from MODIS satellites for Procida due to seawater spectral signal contamination inside each 1 x 1 km MODIS pixel, LST has been estimated from satellite images acquired by OLI and TIRS sensors aboard LANDSAT-8 satellite. LST maps at 30 m spatial resolution were computed using the Plank equation, after estimating brightness temperature and emissivity from LANDSAT-8 satellite spectral bands. A total of 16 cloud free satellite images acquired throughout solar year 2016 have been used to estimate LST, using the “Land Surface Temperature Estimation QGIS Plugin”^60^. A regression analysis has been performed in order to transform LST estimates, describing the skin temperature of earth surface objects, to the located above air temperature, measured by the weather station at the same satellite acquisition time. The analysis allowed to creation of a regional regression model, that has been trained only accounting for LST estimates in the pixel corresponding to meteorological station location. Finally, the regression model has been used to estimate spatial maps of daily air temperature in Celsius degrees for year 2016 from *in-situ* meteorological measurements, accounting for the spatial variability described in LST maps.

### Statistical Analysis

#### Temporal analysis

A Generalized linear additive mixed model (GAMM) was used to assess the relationship between the number of *Ae. albopictus* eggs collected in the 26 ovitraps monitored weekly from April to December 2016 and meteorological variables. GAMM was applied on the series of collected eggs with the following equation:

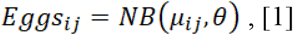

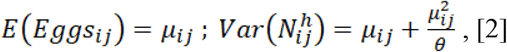

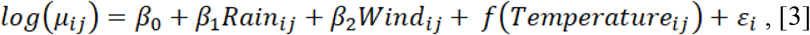

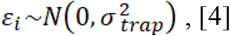

*Eggs_ij_* is the total number of collected eggs at collection week *j* in ovitrap *i* and was assumed to follow a Negative Binomial distribution of mean *μ_ij_* and dispersion parameter *θ* (equation [1] and [2]). A log link function was considered to model *μ_ij_* as a function of independent variables (eq. [3]). *Rain* is the cumulative precipitation during the week of collection, *Wind* is the average wind speed during the week of collection, *f(Temperature)* is the temperature trend modelled by a first order random walk model and *Temperature* is the average temperature during the week of collection. Given that multiple observations were collected from each ovitrap, ovitrap was considered as random effect (ε, eq. [4]). Penalized complexity priors (U=0.05; α = 0.05) were used for the random walk model of the temperature trend, a log-gamma distribution was used for the priors of the log-transformed precision of the random effect trap, while normal distributions of mean 0 and precision 0.001 were used for the priors of β parameters. Finally, all quantitative independent variables were standardized (subtracted their mean value and divided by their standard deviation)^61^. GAMM was fitted in a Bayesian framework using INLA^62^ and R version 3.5.1^63^. Assessment of the statistical assumptions in the model was carried out by autocorrelation function, variogram and graphical inspection of the residuals. Model fit was evaluated by computing the Bayesian p-value for each observation and the conditional predictive ordinate (CPO) using ‘leave one out’ cross validation. 2016 and 2017 meteorological data was used to predict the mean number of eggs in ovitrap for weekly collection and estimate the start and end of the breeding season. The start and end of the season were defined as the week when the cumulative number of eggs collected exceed the 5% and 95% quantile, respectively.

#### Spatial analysis

The geo-referenced field data from the 101 ovitraps monitored in July and September 2016 were identified on the projection system UTM Zone 33N with datum WGS84 (EPSG code 32633), relating to the Italian cartographic system. The vector and raster maps were prepared and visualized on the Open-Source Quantum GIS version 2.18.2 Las Palmas^59^ software and the spatial statistical analysis was undertaken using interpolation by the kriging method on the Open Source R v3.3.2^63^, gstat package^64^. In geo-statistics, the random field (RF) *Z* (*u*) is assumed to be intrinsic second order stationary if the first two moments (i.e. the mean or trend component *m* and the semi-variance *γ*(*h*)) of the two point RF increments exist and are invariant under translation and rotation within a bounded area *D*^65,66^:

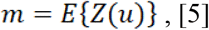

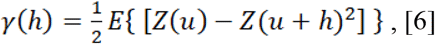

with theoretically infinite points locations *u* (*x*) *D*, and random variables (RV) *Z* (*u*) and *Z* (*u* + *h*) separated by the distance vector *h* (*x*), where *x* represents the spatial coordinates (x_1_, x_2_) ℜ^2^ in our study domain. In ordinary kriging the mean is deemed stationary in the local neighbourhood of locations *u* and unknown, which brings to the following kriging system in matrix notation:

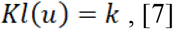

Geostatistical interpolation offers the possibility to use the spatial dependency of the variable under investigation to get, on the basis of field observations, the values of that target variable at any unsampled or unknown location over the whole study area – in our case in the area of the Procida and Vivara islands. Kriging weights:

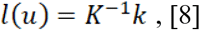

are calculated after the fitting of an experimental variogram through an allowed model variogram, which allows in turn to derive the vector of data to unknown covariance:

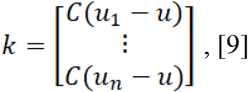

The vector of weights:

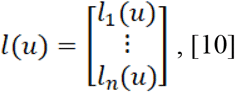

is calculated by solving the kriging system in equation [8], where:

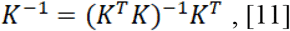

The map of interpolation by ordinary kriging (OK) is calculated for each unknown location *u* of the grid by iterative linear combination of kriging weights with measurements z(u_α_) at sampling locations^65^:

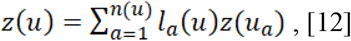

#### Mean distance travelled, Population Survival and Size

Mean distance travelled (MDT) was computed to estimate the dispersal of marked *Ae. albopictus* males from MRR experiments, taking into account the unequal trap densities within each annulus^51,67^. MDT is independent of the position of the traps or size of study area^68–70^, and is defined by the following equation:

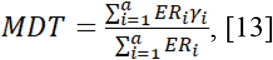

where, *a* is the number of annuli (*a*=4), *γ* is the median distance of each annulus, ER is the number of recaptures that would be expected if trap density was constant within annulus. See supplementary S1 for further details.

Based on MDT results, which provide an estimate of the area to consider in the estimation of the population size, a Generalized Linear Model (GLM) with Binomial distribution was used to estimate the population survival rate of marked *Ae. albopictus* males and the population size of the wild population^49^. GLM was applied on the series of collected marked males out of the total number of collected male mosquitoes with the following equation:

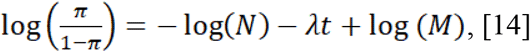

Where N is the population size, M is the number of mosquitoes released, λ is the rate of survival function and t is the time (days) between release and recapture of mark mosquitoes as in Cianci et al. (2013). See supplementary Methods S2 online for further details.

Moreover, the Fisher-Ford’s method modified for low recapture rate^49^ was also applied to provide a second estimate of population size. The Fisher-Ford’s equation is the following:

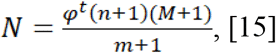

where N is the population size, ϕ is the marked male survival function, t is the time between release and capture, n is the number of both marked and unmarked mosquitoes captured, m is the number of marked mosquitoes recaptured and M is the number of marked mosquitoes released. The confidence intervals related to the estimates are calculated with the method of percentile bootstrap^71^ based on 1000 bootstrap replicates at 95% level.

## Acknowledgements

This work was supported by two grants “Finanziamento Straordinario del Rettore dell’Università degli Studi di Napoli Federico II” to MS in 2016 and 2017, by a grant from Comitato di Gestione della Riserva Naturale Statale Isola di Vivara to MS in 2018 and by EU funded projects H2020-MSCA-NIGHT-2018 and H2020-MSCA-NIGHT-2020. We are deeply grateful for the invaluable help of the Procida Major Raimondo Ambrosino and the municipal counsellors Rossella Lauro and Titta Lubrano. We are very grateful to the Procida volounteers Davide Zeccolella, Luigi “Corecane” D’Orio, Cesare Buoninconti, Amedeo Schiano, Michele Meglio, Alberto Salvemini, Marilena Scotto D’Apollonia, Max Noviello, Anna and Antonio Amalfitano, Emanuela Coppola, Biagio and Isa Coppola, Angela and Pasquale Lubrano, Giulia and Angelo Salvemini, Claudia Riccio and Antonietta Pagano, who shared their time to support this project. We thank Franco e Maria Costagliola and Pina Cuccurullo for the hosting of many students and researchers in their houses during the years of the project. We thank Nella Scotto for the assistance during the surveys on the island. We greatly thank Proff. Luciano Gaudio and Serena Aceto for their encouragement and support during these years of study on the island and for critical reading of the manuscript. We thank students of the University of Naples Federico II involved in this project: Brunella Bozzi, Angela Meccariello, Rita Colonna, Antonia Fiore, Claudia Ascione, Daniela Carannante and Antonio Marino. We thank all the Procida citizens and tourist accommodations (Hotel Riviera, Hotel La Torre, Hotel Savoia, Camping Punta Serra, Camping Vivara) that granted us access to their private properties for mosquito monitoring activities. We thank Pasquale Raicaldo for his support in reporting on national newspaper about our project progresses on the Island. We thank the Comitato di Gestione Isola di Vivara for the authorization to access the reserve. We are deeply grateful to Rui Cardoso Pereira, Jeremy Bouyer, Kostas Bourtzis, Marc Vreysen and Jorge Hendrichs of the Joint FAO/IAEA Division of Nuclear Techniques in Food and Agriculture, Wien, Austria, for their help and support. We thank Jeremy Bouyer, Luciano Gaudio and Serena Aceto for their critical reading of the manuscript.

## Author contributions

M.S., B.C., A.d.T. and R.B. conceived and designed the experiments. M.S. coordinated the study and the logistic and administrative interactions with the Procida Municipality. V.P. coordinated the volunteers involved in the ovitrap monitoring for the temporal analysis. C.D. coordinated and performed the ovitrap monitoring on Vivara island. M.S., B.C., V.P., M.V., C.D., L.I., P.P., L.T., C.R., A.L., G.SO., M.M.P., M.F. and F.P. participated in the field entomological surveys and laboratory operations. P.P. and M.S. collected eggs and larvae to produce *Ae. albopictus* sterile males. A.P. and R.B. produced the sterile males for the MRR experiments. M.S., B.C., M.V., P.P., L.T., A.L., G.SO., M.M.P., M.F. and F.P. performed the MRR experiments. M.S., G.L., B.C., M.M., F.F., C.V., R.R., and R.B. analysed and interpreted the data. M.S., B.C., G.L. and A.d.T. wrote the paper with input of M.M., R.B., R.R., C.V. and G.SA. All authors read and approved the final manuscript. G.SO., M.M.P., M.F., and F.P. were students from Department of Biology (University of Naples Federico II), participating to this research project as volunteers, during the period of their experimental thesis in Genetics (Tutor Prof. G. Saccone). P.P. is a Biology PhD student (University of Naples Federico II), participating to this research project as a volunteer, during the period of his experimental activities in Genetics (Tutor Prof. G. Saccone).

## Data availability

All data generated or analysed during this study are included in this published article (and its Supplementary Information files).

