## Supplementary figures S1-S8 and supplementary Methods S1-S2 for "*Aedes albopictus* bionomics in Procida Island, a promising Mediterranean site for the assessment of innovative and community-based integrated pest management methods"

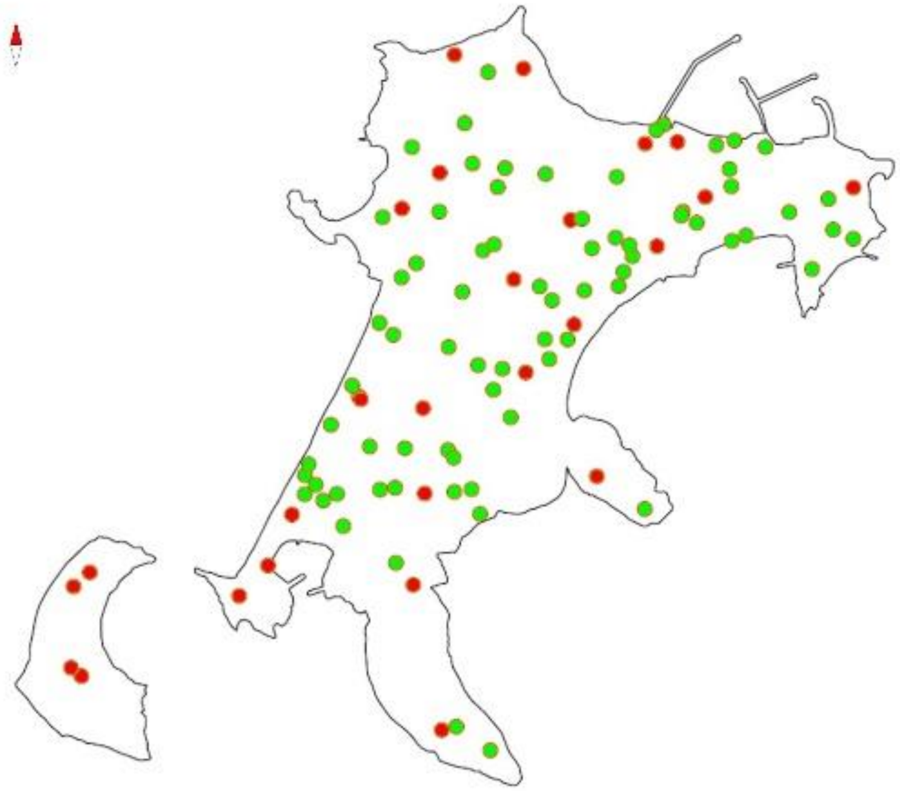

**Figure S1. Position of the ovitraps on Procida Island.** The position of the 101 ovitraps utilized for temporal (red) and spatial (red and green) analyses are reported.

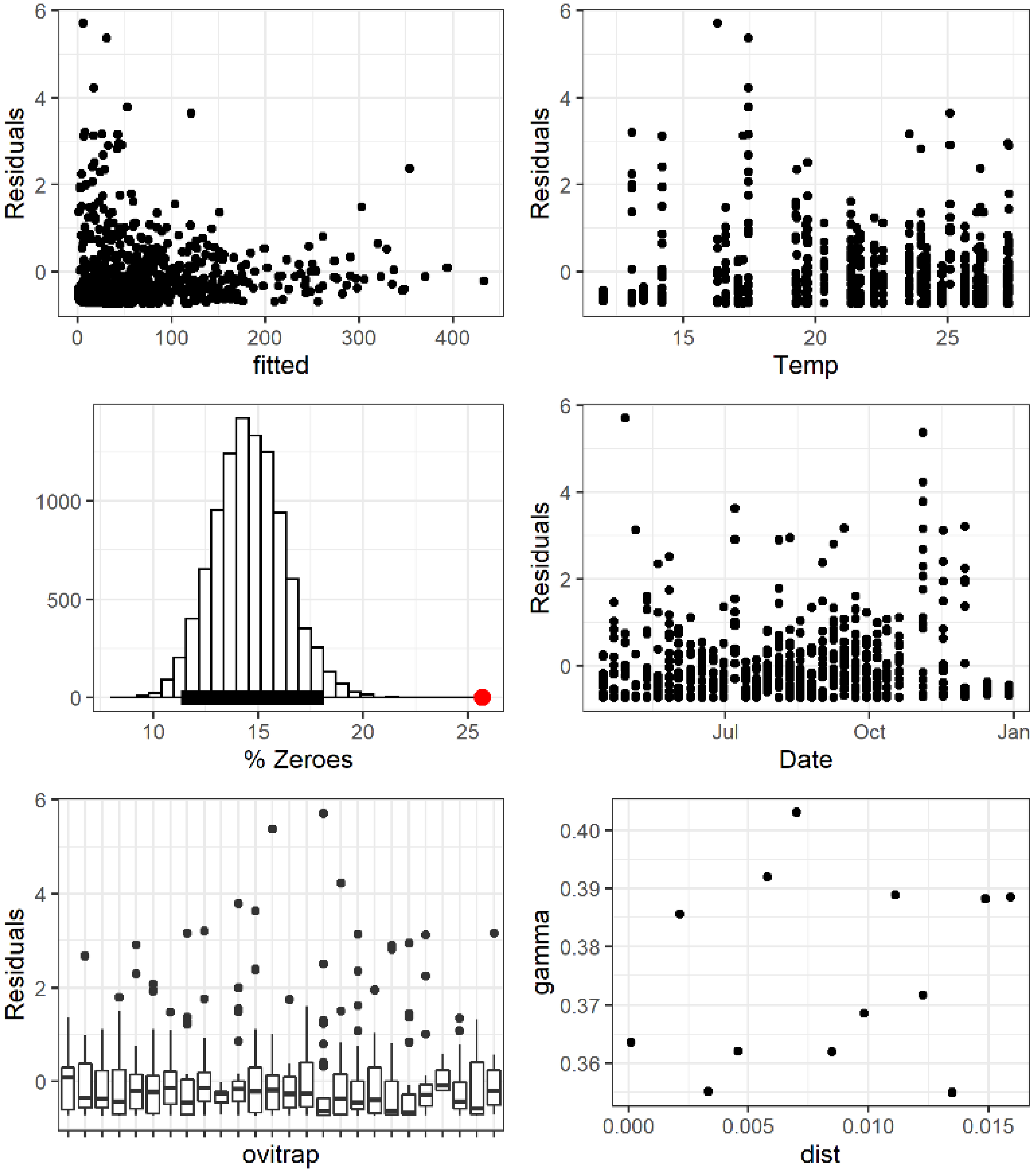


**Figure S2. GAMM model validation.** Upper right panel: Pearson's residuals versus fitted values. Upper left panel: Pearson's residuals versus temperature. Middle right panel: Histogram of % of zeroes obtained by simulating 10000 databases, the red dot represents the observed % of zeroes. Middle left panel: Pearson's residuals versus fitted date of collection. Lower right: Pearson's residuals versus ovitraps. Lower left: Variogram of Pearson's residuals. Autocorrelation function of each ovitrap time series did not show serious violation of independence.

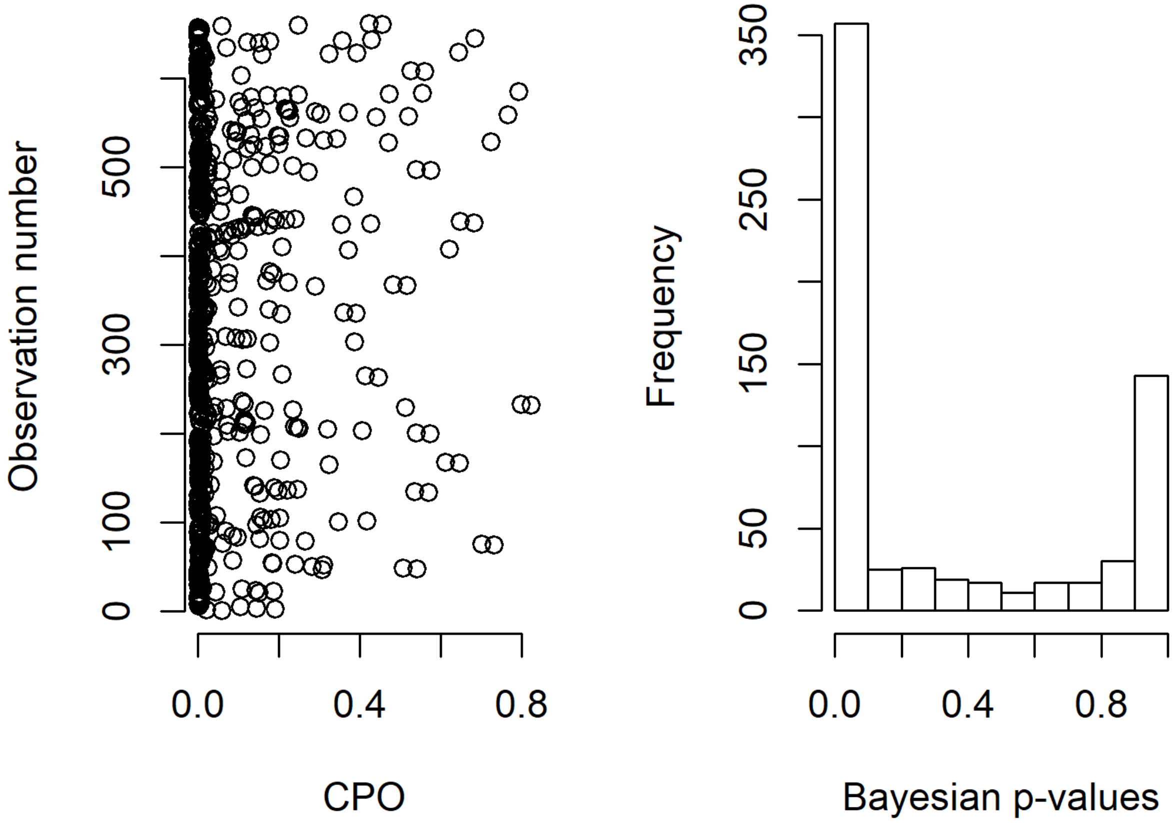


**Figure S3. Assessment of GAMM model fit.** Left panel: On the x axis the conditional predictive ordinate (CPO) which represents the posterior probability of observing that observation when the model is fit using all data except that one. On the y-axis the observation. Right panel: the frequency distribution of the probability of a new value to be lower than the actual observed value (Bayesian p-value).

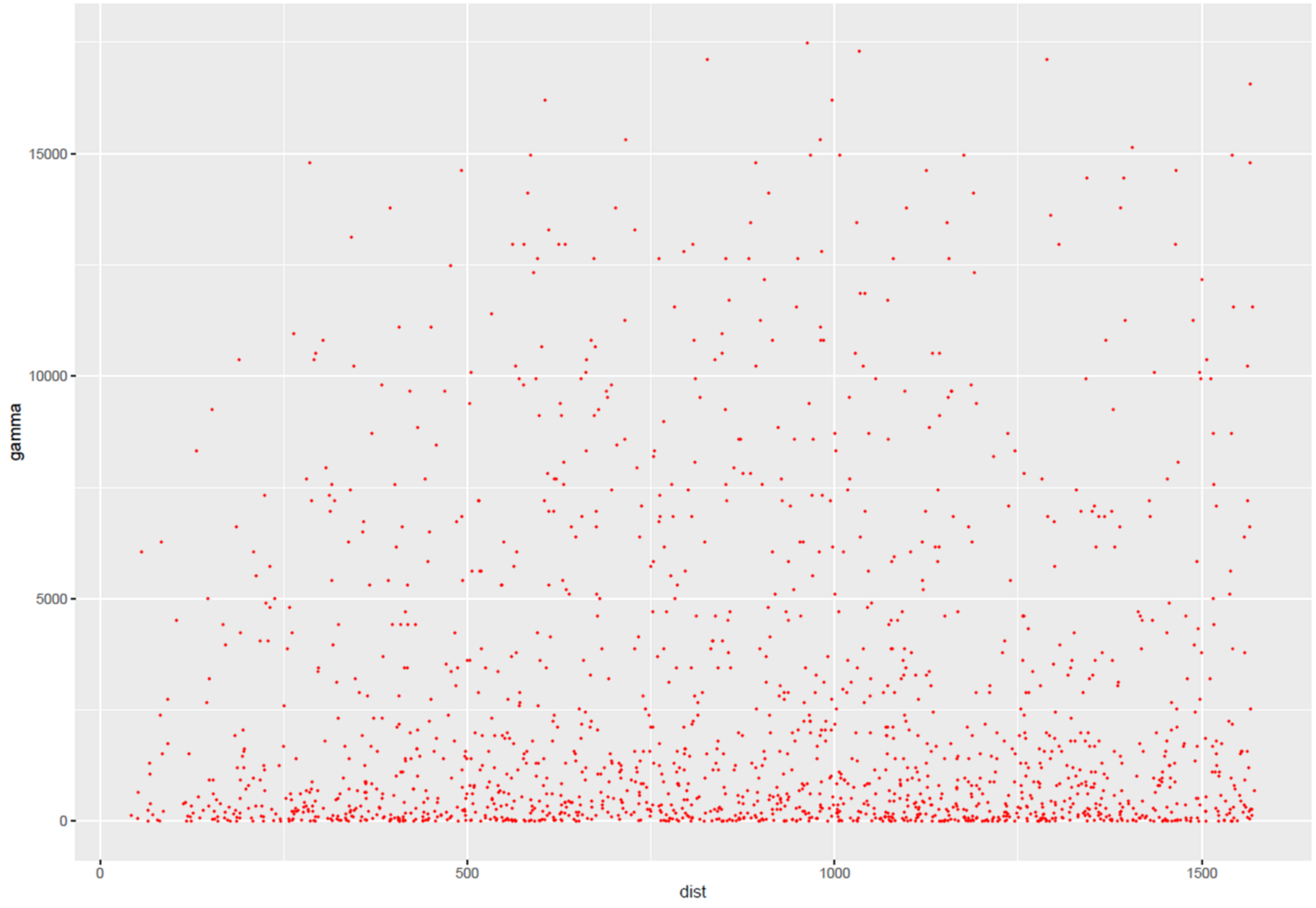


**Figure S4. Example of cloud variogram for week 29.** Each point on the plot represents a couple of two locations separated by a distance vector in 2D spatial domain (x axis) and having a semivariance value reported on y axis.

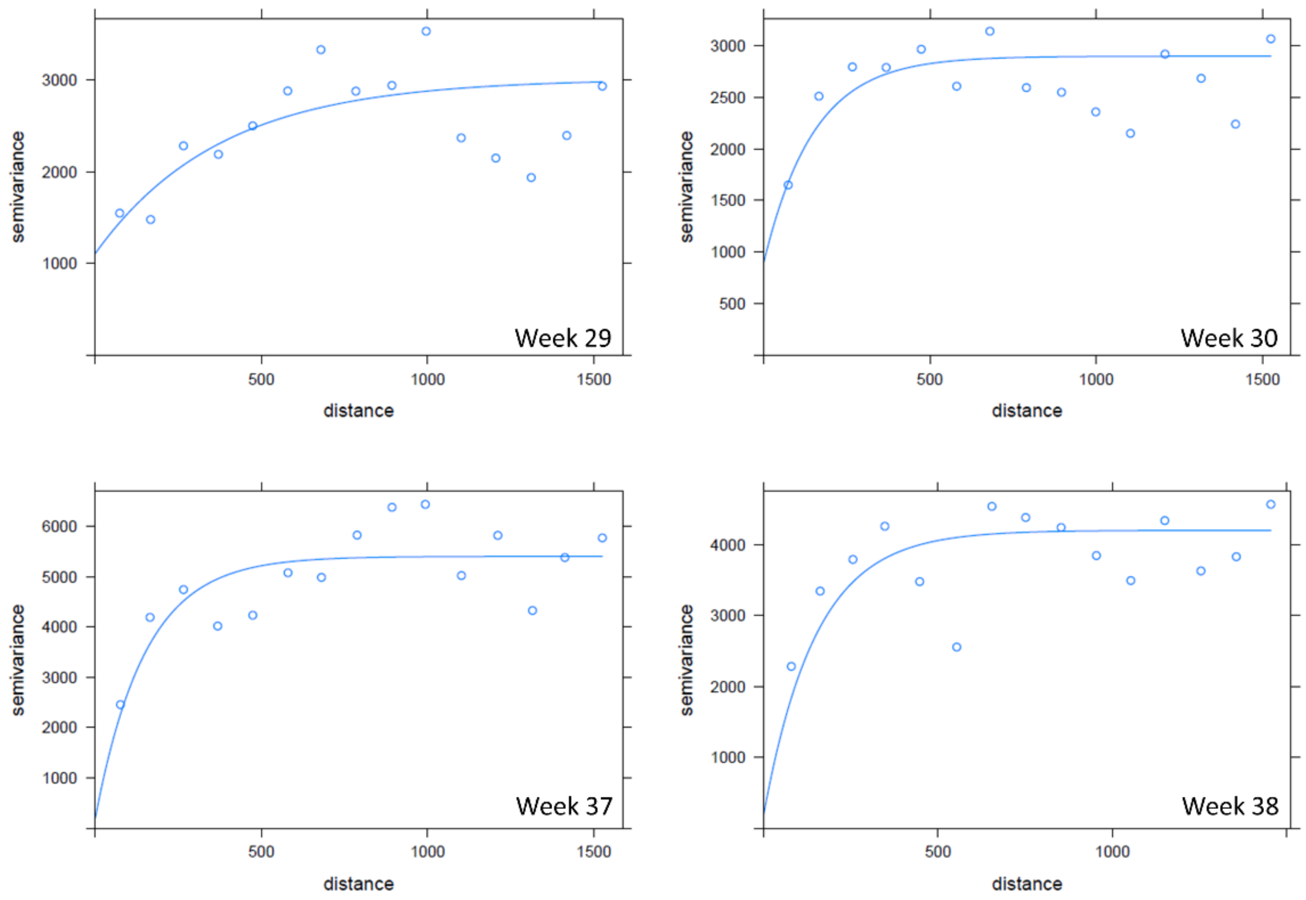


**Figure S5. Experimental semivariograms of total eggs/ovitrap/week in Procida.** Each point represents the average value of semivariance of couple of locations belonging to the same lag. This is called semivariogram and it is the main tool in geostatistics to discover the existence of spatial structure in the data. It is used to inform the interpolation by kriging.

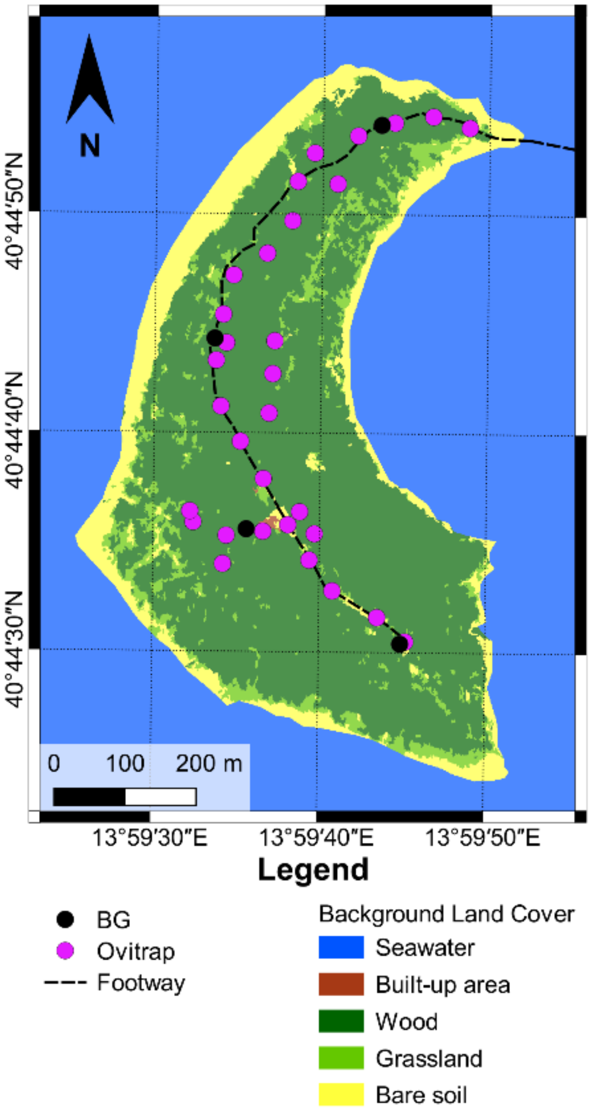


**Figure S6. Positions of ovitraps and BG-traps on Vivara Island.** Purple circles represent the position of the ovitraps. Black circles represent the position of BG-sentinel traps.

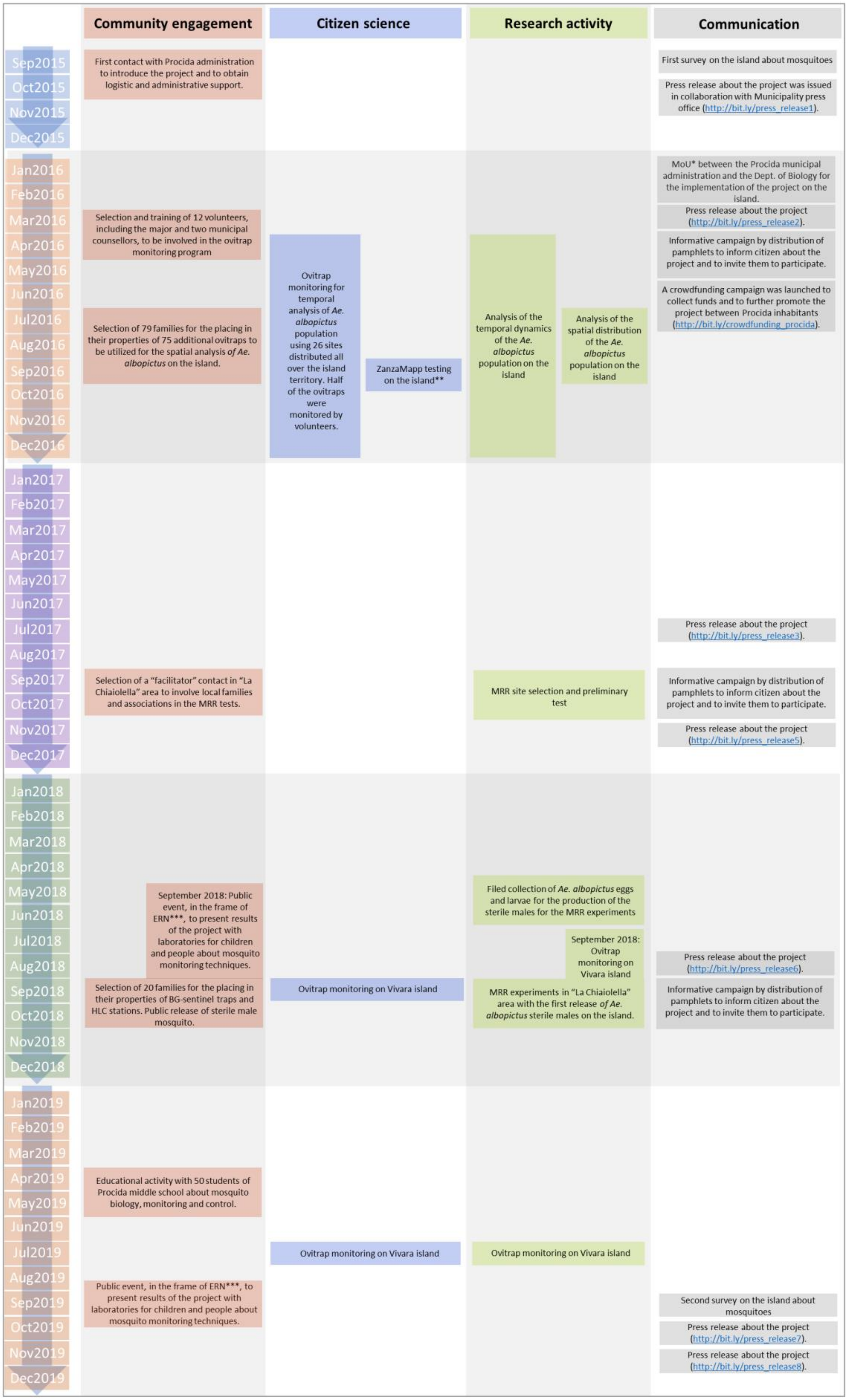

Community engagement
Citizen science
Research activity
Communication
Sep2015
Oct2015
Nov2015
Dec2015
First contact with Procida administration to introduce the project and to obtain logistic and administrative support.
First survey on the island about mosquitoes
Press release about the project was issued in collaboration with Municipality press office (http://bit.ly/press_release1).
Jan2016
Feb2016
Mar2016
Apr2016
May2016
Jun2016
Jul2016
Aug2016
Sep2016
Oct2016
Nov2016
Dec2016
Selection and training of 12 volunteers, including the major and two municipal counsellors, to be involved in the ovitrap monitoring program
Selection of 79 families for the placing in their properties of 75 additional ovitraps to be utilized for the spatial analysis of Ae. albopictus on the island.
Ovitrap monitoring for temporal analysis of Ae. albopictus population using 26 sites distributed all over the island territory. Half of the ovitraps were monitored by volunteers.
ZanzaMapp testing on the island**
Analysis of the temporal dynamics of the Ae. albopictus population on the island
Analysis of the spatial distribution of the Ae. albopictus population on the island
MoU* between the Procida municipal administration and the Dept. of Biology for the implementation of the project on the island.
Press release about the project (http://bit.ly/press_release2).
Informative campaign by distribution of pamphlets to inform citizen about the project and to invite them to participate.
A crowdfunding campaign was launched to collect funds and to further promote the project between Procida inhabitants (http://bit.ly/crowdfunding_procida).
Jan2017
Feb2017
Mar2017
Apr2017
May2017
Jun2017
Jul2017
Aug2017
Sep2017
Oct2017
Nov2017
Dec2017
Press release about the project (http://bit.ly/press_release3).
Selection of a "facilitator" contact in "La Chiaiolella" area to involve local families and associations in the MRR tests.
MRR site selection and preliminary test
Informative campaign by distribution of pamphlets to inform citizen about the project and to invite them to participate.
Press release about the project (http://bit.ly/press_release5).
Jan2018
Feb2018
Mar2018
Apr2018
May2018
Jun2018
Jul2018
Aug2018
Sep2018
Oct2018
Nov2018
Dec2018
September 2018: Public event, in the frame of ERN***, to present results of the project with laboratories for children and people about mosquito monitoring techniques.
Filed collection of Ae. albopictus eggs and larvae for the production of the sterile males for the MRR experiments
September 2018: Ovitrap monitoring on Vivara island
Press release about the project (http://bit.ly/press_release6).
Selection of 20 families for the placing in their properties of BG-sentinel traps and HLC stations. Public release of sterile male mosquito.
Ovitrap monitoring on Vivara island
MRR experiments in "La Chiaiolella" area with the first release of Ae. albopictus sterile males on the island.
Informative campaign by distribution of pamphlets to inform citizen about the project and to invite them to participate.
Jan2019
Feb2019
Mar2019
Apr2019
May2019
Jun2019
Jul2019
Aug2019
Sep2019
Oct2019
Nov2019
Dec2019
Educational activity with 50 students of Procida middle school about mosquito biology, monitoring and control.
Ovitrap monitoring on Vivara island
Ovitrap monitoring on Vivara island
Public event, in the frame of ERN***, to present results of the project with laboratories for children and people about mosquito monitoring techniques.
Second survey on the island about mosquitoes
Press release about the project (http://bit.ly/press_release7).
Press release about the project (http://bit.ly/press_release8).

**Figure S7. Workflow of the community engagement approach utilized during the 4 years of activities in Procida island.** (*) MoU = Memorandum of Understanding. (**) ZanzaMapp is a mobile app for mosquito monitoring (https://www.zanzamapp.it/)[72] that was tested on Procida island during September 2016. The paper which describes the entomological validation of the data collected by citizens on the islands is in preparation (Caputo et al., 2020 in prep.). (***) ERN = European Research Night. The public activities were organized on Procida island in the frame of the MEETmeTONIGHT (http://www.meetmetonight.it/) funded by EU.

**Ovitrap monitoring for temporal analysis April 2016-December 2016**

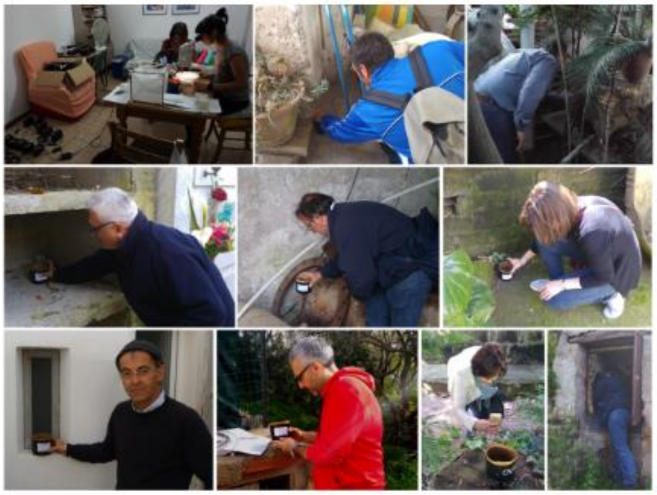

**Ovitrap monitoring for spatial analysis July 2016-September 2016**

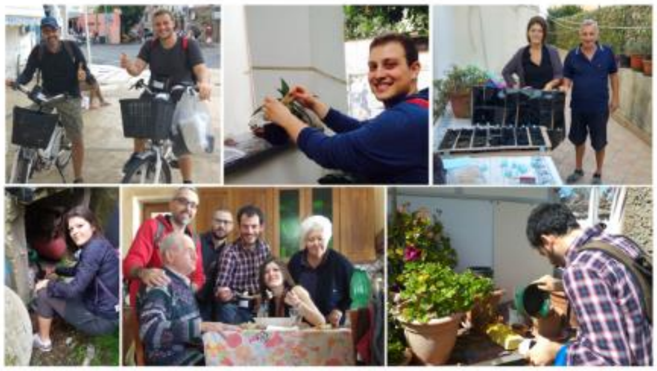

**Mark-release-recapture experiments September 2018**

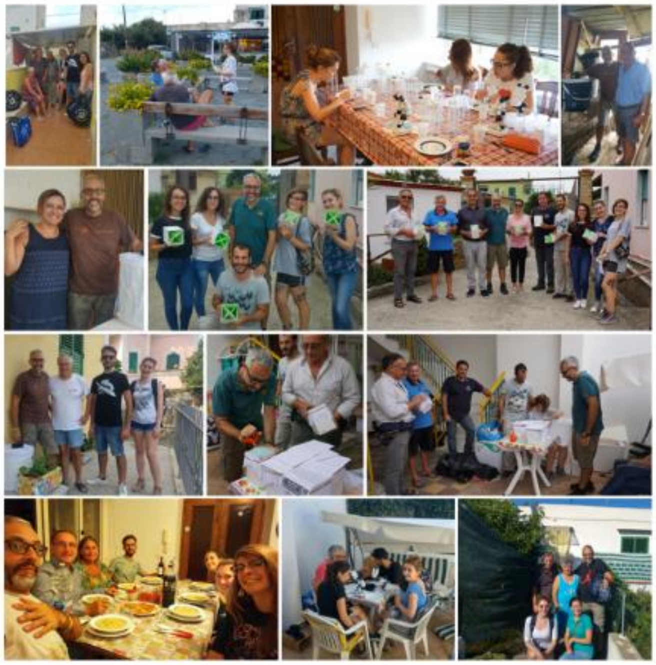

**Figure S8. Volunteers involved during the three phases of the research program on Procida and Vivara Islands.** All the people (co-authors of the present manuscript and Procida volunteers) present in this figure gave their consent to the publication of the images.

**Supplementary Methods S1**

The equation [14] obtained from this equation:

$$\log\left(\frac{\pi}{1-\pi}\right) = -\log(N) - \log\left[S_m(t)\right] + \log(M), \text{ [16]}$$

where π is the population fraction of marked mosquitoes, N is the population size, M is the number of mosquitoes released and $S_m(t)$ is a survival function. Suppose that survival function is an exponential distribution:

$$S_m(t) = e^{-\lambda t} \text{ [17]},$$

the equation [14] become:

$$\log\left(\frac{\pi}{1-\pi}\right) = -\log(N) - e^{\beta_0}t + \log(M),$$

*N* can be estimated from the intercept of the model,

$$\alpha = -\log(N); \text{ [18]}$$

$$\hat{N} = \exp(-\hat{\alpha}). \text{ [19]}$$

The approximate 95% confidence interval has been calculated by:

$$\exp[-\alpha - zSE(\hat{\alpha})] < N < \exp[-\alpha + zSE(\hat{\alpha})], \text{ [20]}$$

Where *z* is a standard normal distribution [49].

**Supplementary Methods S2**

$$ER = \frac{N^{\circ}\ of\ recaptured\ in\ each\ annulus}{N^{\circ} traps\ in\ each\ annulus} * CF. \quad [21]$$

CF is a correction factor to account for differences in trap densities among annuli

$$CF = \frac{\pi(R^2 - r^2)}{\pi R^2} * \text{N° of traps in the study areas.} \quad [22]$$

The Mean distance Travelled values was calculated for the first release and for the second release. The Flight Ranges (FRs) were obtained from the linear regression of the cumulative number of expected recapture (ERs) from each annulus on the $\log_{10}$ (distance) (Marini et al., 2010).

**Table S1. Data and coordinates for the temporal, spatial and MRR analyses.**

**Table S2. Pearson correlation coefficients between ovitrap egg numbers in spatial analysis, terrain parameters derived from digital terrain model (dtm) and sea distance.**
